# Protein Sequence Design with a Learned Potential

**DOI:** 10.1101/2020.01.06.895466

**Authors:** Namrata Anand-Achim, Raphael R. Eguchi, Irimpan I. Mathews, Carla P. Perez, Alexander Derry, Russ B. Altman, Po-Ssu Huang

## Abstract

The task of protein sequence design is central to nearly all rational protein engineering problems, and enormous effort has gone into the development of energy functions to guide design. We investigate the capability of a deep neural network model to automate design of sequences onto protein backbones, having learned directly from crystal structure data and without any human-specified priors. The model generalizes to native topologies not seen during training, producing experimentally stable designs. We evaluate the generalizability of our method to a *de novo* TIM-barrel scaffold. The model produces novel sequences, and high-resolution crystal structures of two designs show excellent agreement with the *in silico* models. Our findings demonstrate the tractability of an entirely learned method for protein sequence design.

---

Computational protein design has emerged as a powerful tool for rational protein design, enabling significant achievements in the engineering of therapeutics [1, 2, 3], biosensors [4, 5, 6], enzymes [7, 8], and more [9, 10, 11]. Key to such successes are robust sequence design methods that minimize the folded-state energy of a pre-specified backbone conformation, which can either be derived from existing structures or generated *de novo.* This difficult task [12] is often described as the “inverse” of protein folding—given a protein backbone, design a sequence that folds into that conformation. Current approaches for fixed-backbone design commonly involve specifying an energy function and sampling sequence space to find a minimum-energy configuration [13, 14, 15], and enormous effort has gone into the development of carefully modeled and parameterized energy functions to guide design, which continue to be iteratively refined [16, 17].

With the emergence of deep learning systems and their ability to learn patterns from high-dimensional data, it is now possible to build models that learn complex functions of protein sequence and structure, including models for protein backbone generation [18, 19, 20] and protein structure prediction [21, 22]; as a result, we were curious as to whether an entirely learned method could be used to design protein sequences on par with energy function methods. Recent experimentally validated efforts for machine learning-based sequence generation have focused on sequence representation learning, requiring fitting to data from experiments or from known protein families to produce functional designs [23]. We hypothesized that by training a model that conditions on local backbone structure and chemical environment, the network might learn residue-level patterns that allow it to generalize without fine-tuning to new backbones with topologies outside of the training distribution, opening up the possibility for generation of de novo designed sequences with novel structures and functions. Moreover, for rational protein design we are often interested in designing structures, interfaces, or residue interactions with angstrom-level specificity. As such, we explored a method in which the neural network explicitly builds side-chain conformers and evaluates full atom structural models, an approach not reported to date.

Conventional energy functions used in sequence design calculations are often composed of pairwise terms that model inter-atomic interactions. Given the expressivity of deep neural networks, or their ability to approximate a rich class of functions, we predicted that a model conditioned on chemical context could learn higher order (multi-body) interactions relevant for sequence design (e.g., hydrogen bonding networks). Furthermore, most energy functions are highly sensitive to specific atom placement, and as a result designed sequences can be convergent for a given starting backbone conformation. For most native proteins, however, the existence of many structural homologs with low sequence identity suggests that there is a distribution of viable sequences that can adopt a target fold, but the discovery of these sequences given a fixed-backbone reference structure is difficult. We hypothesized that a learned model could operate as a “soft” potential that implicitly captures backbone flexibility, producing diverse sequences for a fixed protein backbone.

In this study, we explore an approach for sequence design guided only by a neural network that explicitly models side-chain conformers in a structure-based context (Fig. 1A), and we assess its generalization to unseen native topologies and to a *de novo* protein backbone.

**Figure 1:**
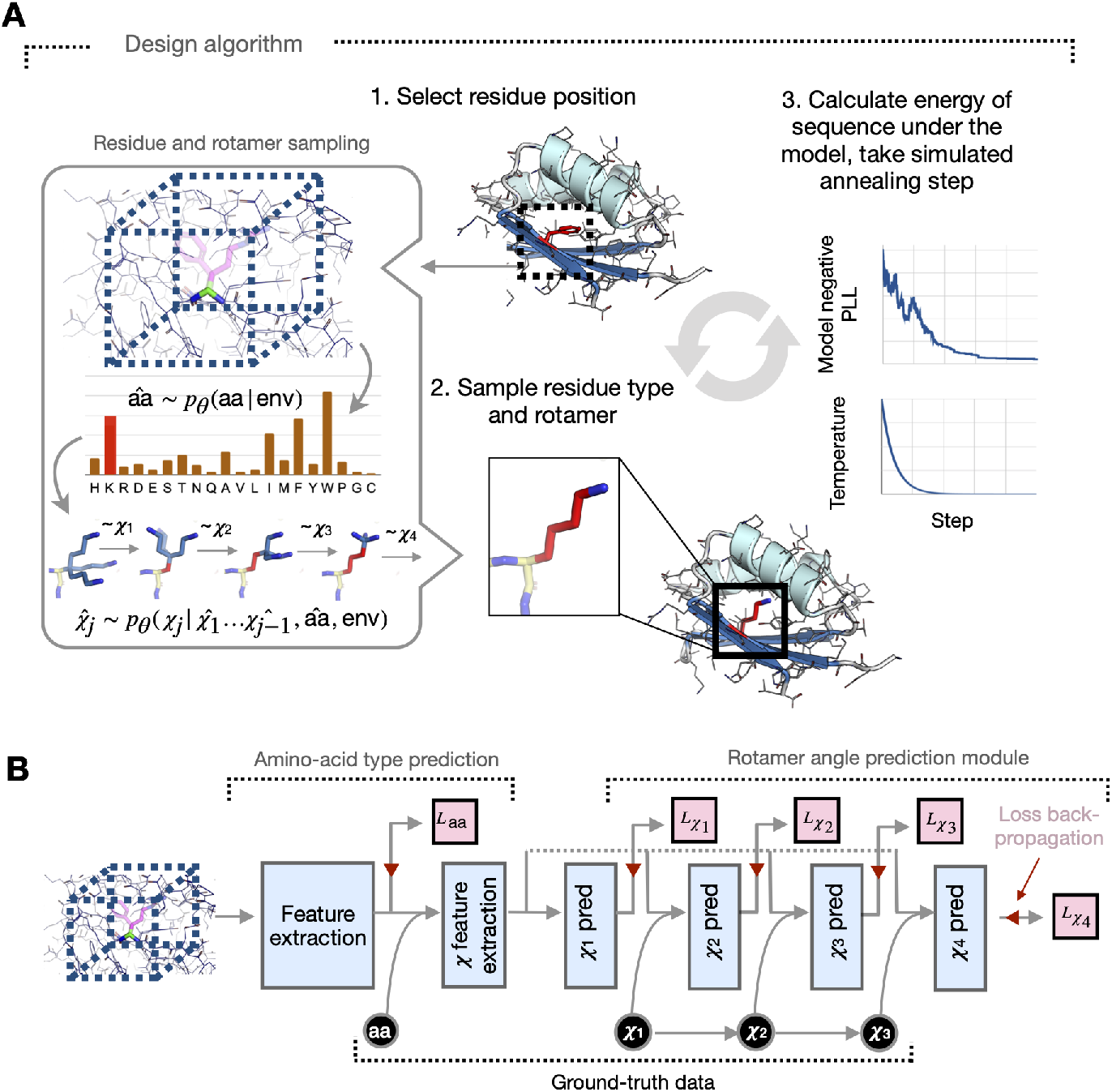
Fully learned sequence and rotamer design onto fixed protein backbones. **A)** Sequences are designed onto fixed protein backbones by (1) iteratively selecting a candidate residue position, (2) using a neural network model to sample amino-acid type and conformation, and (3) optimizing the model “energy” (negative pseudo-log-likelihood of the sequence) via simulated annealing. (Inset, left) Given the local chemical environment around a residue position (box, dashed, not to scale), residue type and rotamer angles are sampled from network-predicted distributions. **(B)** The neural network model is trained to predict residue identity and rotamer angles in an autoregressive fashion, conditioning on ground-truth data (black). The trained classifier predicts amino-acid type as well as rotamer angles conditioned on amino-acid type. Cross-entropy loss objectives shown in pink.

We are interested in sampling from the true distribution of n-length sequences of amino acids *Y* ∈ {1… 20}^*n*^ conditioned on a fixed protein backbone. The backbone is fully specified by the positions of each residue’s four *N – C_α_ – C – O* atoms and the C-terminal oxygen atom, whose positions are encoded as 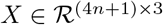; thus, the final conditional distribution we are interested in modeling is:

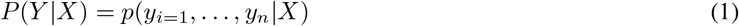

Due to the local nature of the physical interactions within proteins, we can expect that the likelihood of a given side-chain identity and conformation will be dictated by neighboring residues. Defining env_*i*_, as the combination of the backbone atoms *X* and neighboring residues *y*_*NB*(*i*)_, the conditional side-chain distribution at a given position *i* can be factorized sequentially as follows:

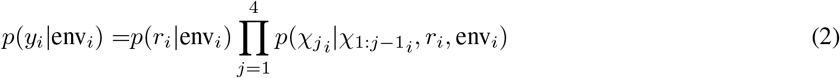

where *r_i_*, ∈ {1… 20} is the amino acid type of residue *i* and *χ*_1_*i*__; *χ*_2_*i*__, *χ*_3_*i*__, *χ*_4_*i*__ ∈ [−180°, 180°] are the torsion angles for the side-chain.

We train a deep neural network **conditional model** *θ* to *learn* these conditional distributions from data. Conditioning on the local environment, the network autoregressively predicts distributions over residue types *p_θ_* (*r_i_*, |env_*i*_) and rotamer angles 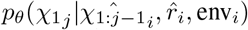 (Fig. 1B). Our design algorithm involves iteratively sampling side-chains conditioned on their local chemical environments from these network-predicted distributions. We can approximate the joint probability of a sequence *P*(*Y |X*) by the pseudo-log-likelihood (PLL) [24] of the sequence under the model

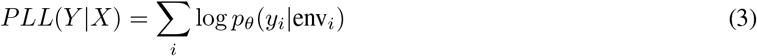

which we optimize in order to find high-likelihood sequences under the model. Over the course of design, the algorithm builds full-atom structural models that can be evaluated using established structure quality metrics.

We use a 3D convolutional neural network as our classifier *θ*, training the model on X-ray crystal structures of CATH 4.2 S95 domains [31, 32, 33], with train and test set domains separated at the topology level. For the amino acid type prediction task, our conditional model achieves a 57.3% test set accuracy, either outperforming [34] or matching [35, 36, 37] previously reported machine learning models for the same task. The predictions of the network correspond well with biochemically justified substitutability of the amino acids (Fig. S1A-C); this type of learned degeneracy is a necessary feature for design, as we expect proteins to be structurally robust to mutations at many residue positions. The residue type-conditioned rotamer prediction module by construction learns the backbone-dependent joint rotamer angle distribution *p*(*χ*_1_*i*__, *χ*_2_*i*__, *χ*_3_*i*__, *χ*_4_*i*__|*X, r_i_*). The network-learned independent χ distributions match empirical residue-specific rotamer distributions, which rotamer libraries typically seek to capture (Fig. S2).

We sought to assess the degree of generalization of the algorithm to native backbones from the test set, which have CATH defined *topologies* not seen by the model during training. We selected four test case backbones that span the major CATH *classes—*all alpha, alpha-beta, and all beta (Fig. 2A). To validate the entirely learned approach, we challenged the model to fully redesign sequences given starting backbones. If the model has generalized, it should be able to recover native rotamers and sequence to a degree during design, as well as design key structural and biochemical elements typically seen in folded proteins.

**Figure 2:**
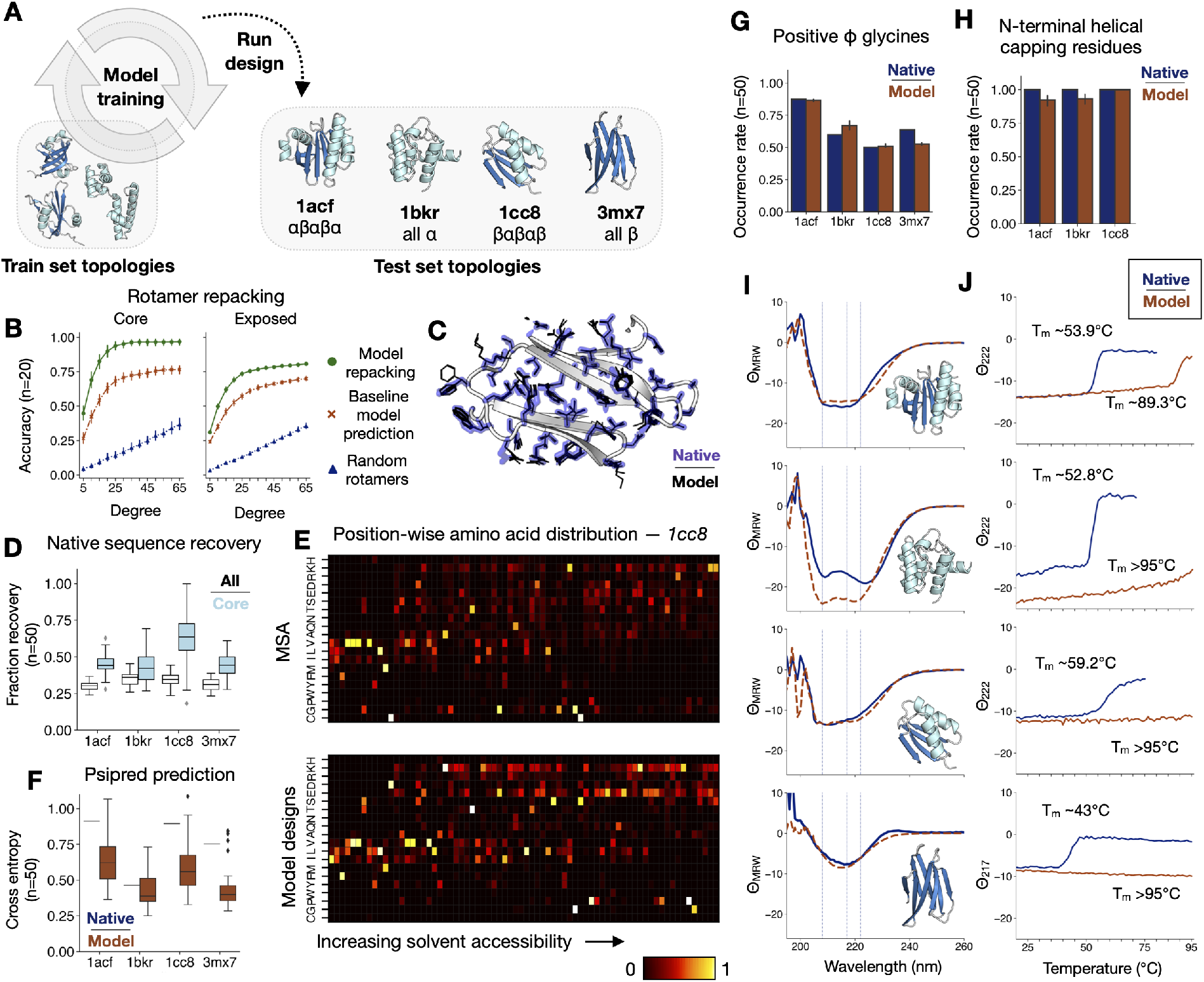
Generalization of model design to unseen topologies. **(A)** The trained model is used to either repack rotamers or design entirely new sequences onto unseen test set structures with non-train-set CATH topologies. (**B** and **C**) Model-guided rotamer recovery for native test cases. **(B)** Rotamer repacking accuracy for buried core residues versus solvent-exposed residues as a function of degree cutoff **(C)** 5 models superimposed with side chains shown as black lines compared to the native conformation shown in purple outline for test case *3mx7.* (**D-H**) Performance of sequence design onto test case backbones. **(D)** Native sequence recovery rate across 50 designs for all residues vs. buried core residues. **(E)** Position-wise amino acid distributions for test case *1cc8.* Columns are normalized. (Top) Native sequence and aligned homologous sequences from MSA (*n* = 670); (Bottom) Model designs (*n* = 50). **(F)** Cross-entropy of Psipred secondary structure prediction from sequence with respect to DSSP assignments [25, 26, 27, 28, 29, 30]. **(G)** Fraction occurrence of glycines at positive *φ* backbone positions across test cases. **(H)** Fraction occurrence of N-terminal helical capping residues across designs for test cases with capping positions. (**I** and **J**) Far UV circular dichroism (CD) spectroscopy data for selected test case designs. **(I)** Mean residue ellipticity *Θ_MRW_* (10^3^deg cm^2^ dmol^−1^) for CD wavelength scans at 20°C for the native structure (blue, dashed) vs. the model design (orange, solid). **(J)** Thermal melting curves monitoring *Θ_MRW_* (10^3^deg cm^2^ dmol^−1^) at 222nm or 217nm for *3mx7.*

Given the native sequences for the test cases, the model recovers native rotamers with high accuracy for the test case backbones across 5 design trajectories each (72.6% accurate within 20 degrees) and with higher accuracy in the hydrophobic core regions of the protein (90.0% accurate within 20 degrees) (Fig. 2B-C, S3A-B); this performance is on par with top benchmarked methods such as Rosetta [38] and another learned method for scoring rotamers [39]. Tasked with designing the sequence and rotamers from scratch (see Methods), the model designs recapitulate between 25-45% of the native sequence in general, with greater overlap in the buried core regions of the protein that are more constrained and therefore might accommodate a limited set of residue types (Fig. 2D, S3C). The model designs are more variable in solvent-exposed regions, akin to sequences homologous to the native structure found through multiple sequence alignment (MSA) (Fig. 2E). Furthermore, the secondary structure prediction accuracy for model-designed sequences are comparable to that of the native sequence (Fig. 2F, S4C), indicating that despite variation, the designed sequences retain local residue patterns that allow for accurate backbone secondary structure prediction.

Model designs tend to have well-packed cores (Fig. S5A-B) and in general, the model-designed sequences do not have hydrophobic residues in solvent-exposed positions, likely due to the abundance of cytosolic protein structures available (Fig. S5C). Additionally, the model designs match the native structure in terms of numbers of side-chain and backbone buried unsatisfied hydrogen bond acceptors and donors (Fig. S5D-F); this indicates that over the course of model design, polar side-chains that are placed in buried regions are adequately supported by backbone hydrogen bonds or by design of other side-chains that support the buried residue.

We also see a number of expected structural features across the test case designs, including placement of glycines at positive *ϕ* backbone positions (Fig. 2G, S4D), N-terminal helical capping residues (Fig. 2H, S4E), universal design of a proline at the cis-peptide position P21 for *1cc8* (Fig. S4F), and, by inspection, polar networks supporting loops and anchoring secondary structure elements.

For each test case backbone, we selected 4 out of 50 designs for further characterization based on ranking by the model PLL and other metrics (Table S1). Importantly, this ranking uses no information about the native sequences. We validated these sequences by computational structure prediction by the Rosetta AbInitio application (Fig. S7, S8). The model designs achieved far better recovery than a 50% randomly perturbed control, suggesting that close recovery of the native backbone is due to features learned by the model and not simply due to sequence overlap with the native.

Given the method’s strong performance under these sequence quality metrics, we sought further confirmation that the model designs would fold *in vivo.* Of the 16 designs tested, 15 expressed well, and 10 appeared well-folded under circular dichroism (CD) wavelength scans (Fig. S9, S10). For each test case, at least 1 of 4 designs appeared folded and had the expected secondary structure signature under CD. For example, the top design for *1bkr* has 208nm and 222nm alpha helical signal as expected for a helical bundle, while all of the 3mx7 designs have clear 217nm beta strand signal, but no alpha signal, consistent with all-beta proteins (Fig. 2I). CD spectra for the top model designs match the native spectra well, and the designs were found to be more thermally stable than the native as well (Fig. 2J, S11). Overall, these results indicate that the machine learning model generalizes to topologies that are strictly unseen by the model during training.

Unlike an analytical energy function, the model is not expected to generalize to structures far from the training distribution (e.g. unfolded, distended, or highly perturbed backbones), as it is trained by conditioning on correct context. However, the model can in fact detect out-of-distribution backbones for a given sequence. Across a set of structures (decoys) generated via fragment sampling during Rosetta AbInitio prediction, the model negative PLL is lowest for decoys with low RMSD to the native backbone (Fig. 3A, S12A). Additionally, in the low backbone RMS regime (< 5Å from the native), the model can further identify decoys with low RMSD side-chain conformations (Fig. 3B, S12B). In essence, though the model is trained to learn sequence dependence on structure, it makes correct predictions about structures given sequence. Notably, the model demonstrates sensitivity to large perturbations, while remaining robust to small perturbations in the backbone (< 2.5Å) and side-chains (< 0.40Å), a desirable property for the design method. To assess whether the model could perform sequence design for *de novo* structures, we tested our method on a Rosetta-generated four-fold symmetric de novo TIM-barrel backbone, a (*βαβα*)_4_ topology consisting of an eight-stranded barrel surrounded by eight alpha helices [40]. Design of *de novo* proteins remains a challenging task as it necessitates generalization to non-native backbones that lie near, but outside the distribution of known structures. Successful *de novo* protein designs often lack homology to any known native sequences despite the fact that *de novo* structures can qualitatively resemble known folds [41, 40, 42]. For a design protocol to perform well on de novo backbones it must therefore supersede simple recapitulation of homologous sequences. The TIM-barrel design case is of particular interest, as about ~ 10% of known enzymes are thought to adopt a TIM-barrel fold [43], making the fold a prime candidate for rational design of enzymes and more generally for substrate binding.

**Figure 3:**
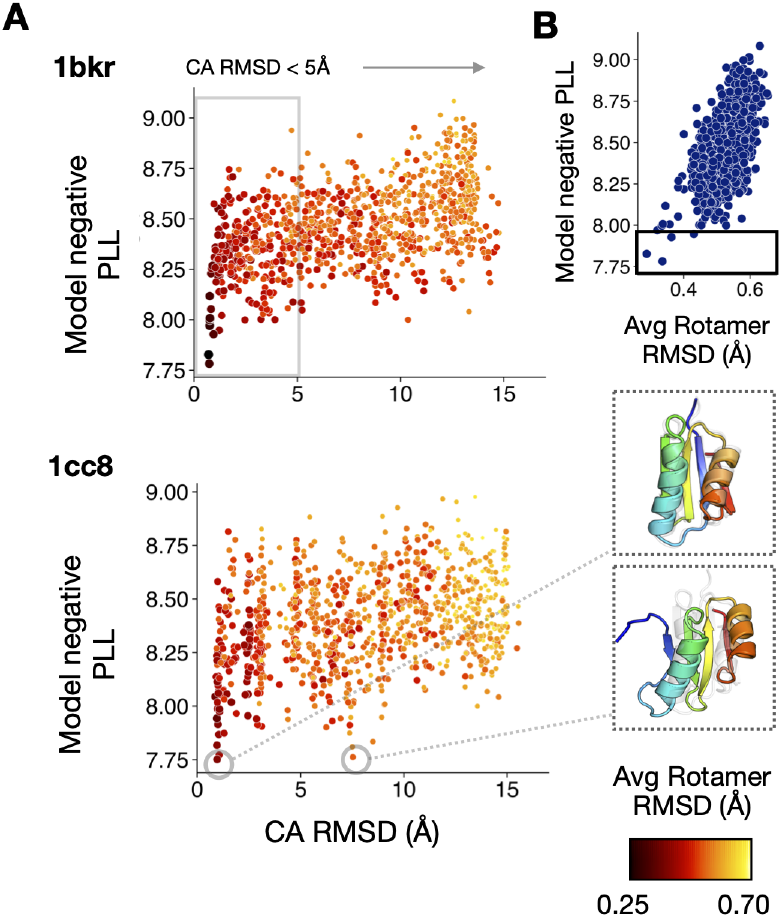
Model captures sequence-structure relationship. (**A-B**) Decoy ranking by model negative pseudo-log-likelihood (PLL) of native sequence. (**A**) Model negative PLL vs. alpha-carbon RMSD (Å) to the native structure for Rosetta AbInitio decoys. Points are colored by average side-chain RMSD to native (Å). In some cases, the model assigns low negative PLL to high RMS backbones; for example, for *1cc8* an alternative pattern of beta strand pairing is shown. (**B**) Model negative PLL of low backbone RMS structures (CA RMSD < 5 Å) vs. average side-chain RMSD (Å). Box highlights low model negative PLL assigned to low side-chain RMSD decoys.

We had the model fully redesign 50 four-fold symmetric sequences for the backbone and selected 8 designs for further characterization based on ranking by the model PLL and other metrics (see Methods), using no information about previously confirmed sequences (Tables S2, S3). A subset of these 8 structures were predicted to fold with low backbone deviation (< 4 Å) into the target backbone by Rosetta (F2C, F4C) and trRosetta (F2C, F4C, F8C) (Fig. S13).

Of these 8 designs, 4 are cooperatively folded proteins, as indicated by circular dichroism (CD) wavelength scans and by the presence of clear two-state transitions in thermal melt curves (Fig. S14, S15). All of the folded proteins designed by our model have higher thermal stability than the initial designs reported in the original study, with melting temperatures for 3 of the 4 ranging between 84-87°C while the fourth protein (F5C) remains folded at 95°C (Fig. S15B). Folded sequences reported in the original study also require manual intervention to introduce a key Asparagine residue required for the structure to cooperatively fold; while automated RosettaDesign is unable to produce foldable sequences, our model produced the sequences with no manual intervention.

We successfully crystallized 2 model designs, F2C (1.46 Å resolution) and F15C (1.9 Å resolution) (Fig. 4A), and the crystal structures validate that the sequences indeed fold into the TIM-barrel structure in excellent agreement with the designed backbone and side-chain rotamers (Fig. 4B, Table S6). Some crystallography artifacts are observed: F2C has a displaced C-terminal helix (Fig. S16A); for two monomers in the asymmetric unit of F15C, one of the the *β-α* loop dislodges to interact with an adjacent monomer (Fig. S16C). Nonetheless, the remaining 3 symmetric replicas are correctly folded. The proteins elute as monomers in size exclusion chromotography (Fig. S14).

**Figure 4:**
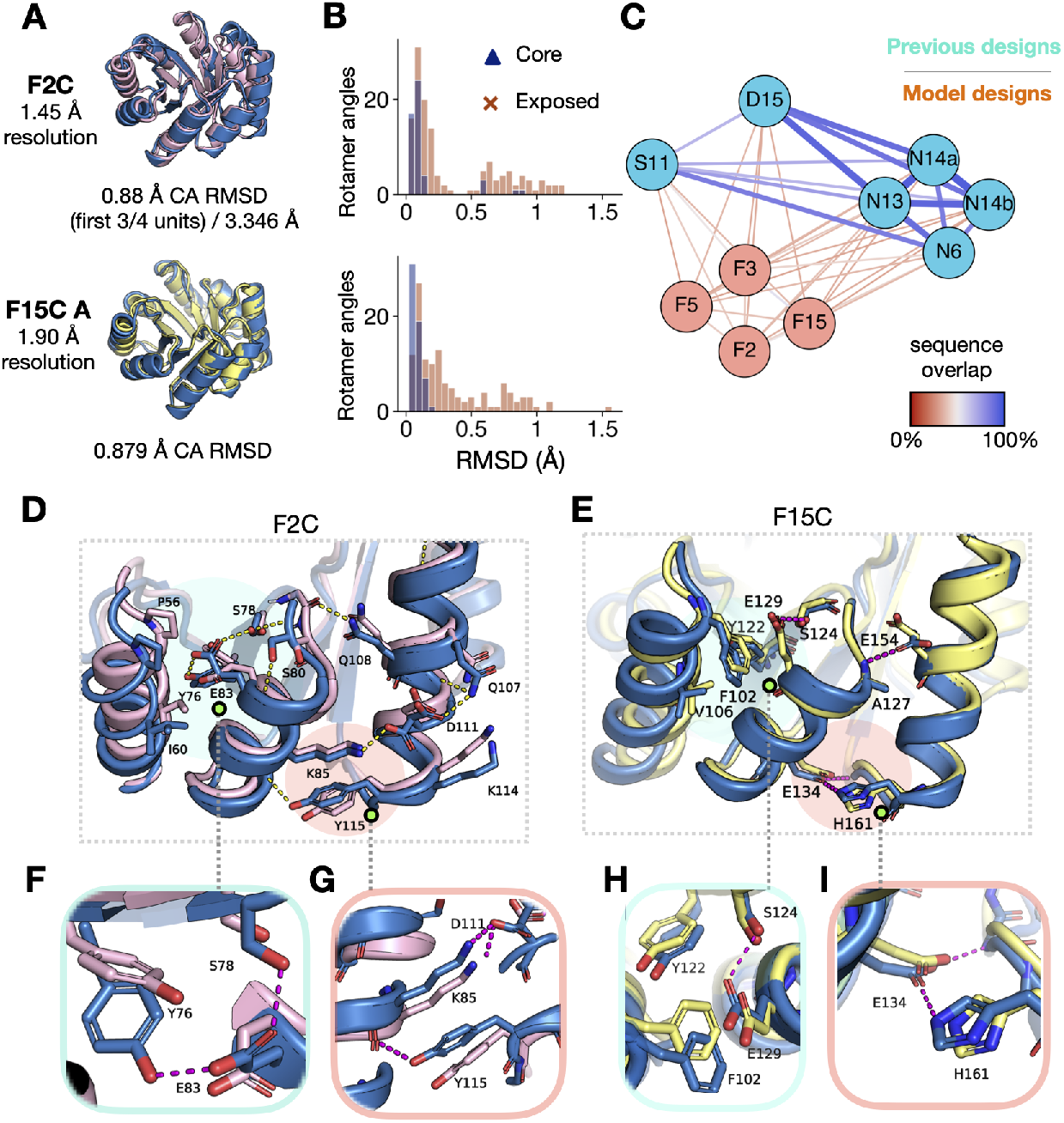
Model discovery of novel sequence features. (**A**) Overlay of crystal structures (blue) with template TIM-barrel backbone for F2C (pink) and F15C (yellow). Alpha-carbon RMSD (Å) given below structures. (**B**) Distribution of rotamer automorphic RMSDs (Å) between crystal structure and template backbone for core and solvent-exposed residues for (top) F2C and (bottom) F15C. (**C**) Sequence overlap between TIM-barrel subunits for model TIM-barrel designs (orange) and previously characterized sequences for the same scaffold (blue), including sTIM-11 *(5bvl,* S11) [40], DeNovoTIM15 (*6wvs*, D15) [44], and NovoTIMs (N6, N13, N14a, N14b) [45]. Edge thickness increases with overlap. (**D-E**) Investigation of sequence features for the symmetric subunit near the top of the barrel (cyan shadow) and the helix interface between symmetric subunits (orange shadow) for (**D**) F2C and (**E**) F15C. Crystal structures shown in blue overlaid with design template (pink – F2C, yellow – F15C). (**F-I**) Closer inspection of novel sequence features designed by the model.

The model designs tend to recapitulate previous core sequences for the region of the protein between the helices and outer part of the barrel (Fig. S17A), as well as for the inner part of the barrel, although some model designs do introduce novel features into these regions (Fig. S17B).

These similarities aside, the model designs are in general more varied than other previously characterized designs, predominantly due to variation in non-buried residue positions (Fig. 4C, S17C, Table S4). Of particular interest are mutations in the interfaces between the helices surrounding the barrel where, remarkably, the design algorithm is able to find novel structural features that have not been discovered by Rosetta and human-guided design methodologies (Fig. 4D-E, S18A-B). For F2C, the model uses an isoleucine to wedge between the long and short helices and forming a Tyr-Glu-Ser polar network that extends to coordinate the loop from the beta sheet to the short helix (Fig. 4F); for F15C, the model has a different solution, designing phenylalanine and valine residues (F10, V14) to pack the long helix against the shorter one near the top of the barrel (Fig. 4H). Particularly interesting are the Tyr-backbone contact and the His-Glu-backbone polar network for F2C and F15C, respectively, that help coordinate the helices across symmetric subunits at the base of the structure; these are the only hydrogen bonding solutions designed at this position among all previous structure-confirmed sequences for this scaffold (Fig. 4G, I).

Our results demonstrate that a design algorithm guided by an entirely learned neural network potential can generate viable sequences for a fixed backbone structure, generalizing to unseen topologies and de *novo* backbones. The method is flexible: the design protocol easily allows for adding position-specific constraints during design, and other neural network models such as graph networks or rotationally equivariant networks, could be used in place of the classifier network presented without fundamentally changing the method. Though presented in a stand-alone manner, in practice this method could be used in tandem with energy function based methods for design, for example by using the model as a proposal distribution, while optimizing an external energy function. We anticipate that this type of approach could be used for design of water-mediated contacts, interfaces, protein-nucleic acid complexes, and ligand binding sites.

Notably, the design algorithm reflects key characteristics of energy functions, such as the ability to (1) accurately determine side-chain conformations, (2) differentiate the hydrophobic interior and polar exterior of the proteins, and (3) design multi-body interactions as reflected by readily produced hydrogen-bonding networks. With classical molecular mechanics force-fields, capturing these effects require terms that accurately describe van der waals, solvation, hydrogen bonding, as well as many other interactions; it also likely requires an independent hydrogen-bonding network search algorithm [46] and discrete side-chain representations from rotamer libraries. None of these are required with this approach. In contrast to energy function development, the model took only hours to train.

In this study, we sought to tackle key challenges with machine-learning based protein sequence design, including zero-shot generalization to new folds and Å-level recovery of target structures. The learned neural network potential is able to guide high-dimensional sampling and optimization of the structure-conditioned sequence distribution. Our results show the practical applicability of an entirely learned method for protein design, and we believe this study demonstrates the potential for machine learning methods to transform current methods in structure-based protein design.

## Methods

### Model training

Our model is a fully convolutional neural network, with 3D convolution layers followed by batch normalization [47] and LeakyReLU activation. We regularize with dropout layers with dropout probability of 10% and with L2 regularization with weight 5 × 10 ^−6^. We train our model using the PyTorch framework, with orthogonal weight initialization [48]. We train with a batch size of 2048 parallelized synchronously across eight NVIDIA v100 GPUs. The momentum of our BatchNorm exponential moving average calculation is set to 0.99. We train the model using the Adam optimizer (β_1_ = 0.5, β_2_ = 0.999) with learning rate 7.5 × 10 ^−5^ [49]. We use the same model architecture and optimization parameters for both the baseline (prediction from backbone alone) and conditional models (Dataset S1).

Our final conditional classifier is an ensemble of four models corresponding to four concurrent checkpoints. Predictions are made by averaging the logits (unnormalized outputs) from each of the four networks. Pre-trained models are available at https://drive.google.com/file/d/1cHoyeI0H_Jo9bqgFH4z0dfx2s9as9Jp1/view?usp=sharing.

### Data

To train our classifier, we used X-ray crystal structure data from the Protein Data Bank (PDB) [31], specifically training on CATH 4.2 S95 domains [32, 33]. We first applied a resolution cutoff of 3.0 Å and eliminated NMR structures from the dataset. We then separated domains into train and test sets based on CATH topology classes, splitting classes into ~ 95% and 5%, respectively (1374 and 78 classes, 53414 and 4372 domains each, see Dataset S1). This ensured that sequence and structural redundancy between the datasets was largely eliminated. During training, we did not excise domains from their respective chains but instead retained the complete context around a domain. When a biological assembly was listed for a structure, we trained on the first provided assembly. This was so that we trained primarily on what are believed to be functional forms of the protein macromolecules, including in some cases hydrophobic protein-protein interfaces that would otherwise appear solvent-exposed.

The input data to our classifier is a 20 × 20 × 20 Å^3^ box centered on the target residue, and the environment around the residue is discretized into voxels of volume 1 Å^3^. We keep all backbone atoms, including the *C_α_* atom of the target residue, and eliminate the *C_β_* atom of the target residue along with all of its other side-chain atoms. We center the box at an approximate *C_β_* position rather than the true *C_β_* position, based on the average offset between the *C_α_* and Cβ positions across the training data. For ease of data loading, we only render the closest 400 atoms to the center of the box.

We omit all hydrogen atoms and water molecules, as well as an array of small molecules and ions that are common in crystal structures and/or possible artifacts (Dataset S1). We train on nitrogen (N), carbon (C), oxygen (O), sulfur (S), and phosphorus (P) atoms only. Ligands are included, except those that contain atoms other than N, C, O, S, and P Bound DNA and RNA are also included. Rarer selenomethionine residues are encoded as methionine residues. For the baseline model, we omit all side-chain atoms while training, so that the model conditions only on backbone atoms. For the conditional model, the input channels include: atom type (N, C, O, S, or P), indicator of backbone (1) or side-chain (0) atom, and one-hot encoded residue type (masked for backbone atoms for the center residue). For the baseline model, the input channels only encode atom type, since all atoms are backbone atoms and we assume no side-chain information is known.

We canonicalize each input residue environment in order to maximize invariance to rotation and translation of the atomic coordinates. For each target residue, we align the N-terminal backbone *N – C_α_* bond to the x-axis. We then rotate the structure so that the normal of the *N – C_α_ – C* plane points in the direction of the positive z-axis. Finally, we center the structure at the effective *C_β_* position. By using this strategy, we not only orient the side-chain atoms relative to the backbone in a consistent manner (in the positive z-direction), but also fix the rotation about the z-axis. We then discretize each input environment and one-hot encode the input by atom type.

### Design algorithm

We would like to sample from the probability distribution of *n*-length sequences of amino acids *Y* ∈ {1… 20}^*n*^ conditioned on a fixed protein backbone configuration. The backbone is specified by the positions of the residues’ non-hydrogen atoms whose positions are encoded as 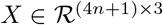; thus, the final conditional distribution we are interested in modeling is:

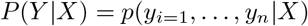

This is a high dimensional structured probability distribution, and we make the following simplifying assumption to make the task of learning this distribution from data more tractable: identity and conformation (rotamer) of each side-chain *y_i_*, is independent of all other side chains *y_j_* conditioned on the identity of the side chains in its neighborhood *y*_*NB*(*i*)_.

This assumption motivates the use of a Conditional Markov Random Field (MRF) to model the target distribution, wherein the nodes of the MRF correspond to the residues and rotamer configurations, edges between nodes indicate the possibility of correlation between the residues or rotamers, and each node is conditionally independent of all other nodes, conditioned on the nodes in its Markov Blanket. More precisely, the input backbone *X* defines a graph structure, where nodes correspond to side chain *y_i_*, and edges exist between pairs of residues (*y_i_ y_j_*) if and only if the corresponding backbone atoms are within some threshold distance of each other. Defining env_*i*_, as the combination of the graph structure *X* and the spatial neighborhood *y*_*NB*(*i*)_, we see that the conditional distribution of residue and rotamers at a single position i can be factorized sequentially as follows:

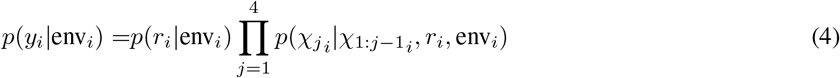

where *r_i_*, ∈ {1… 20} is the amino acid type of residue *i* and *χ*_1_*i*__, *χ*_2_*i*__ *χ*_3_*i*__, *χ*_4_*i*__, ∈ [−180°, 180°] are the torsion angles for the side-chain.

We train a deep neural network **conditional model** *θ* in an autoregressive manner to *learn* these conditional distributions from data (Fig. 1A). We also train a **baseline model** that has only backbone atoms as an input, as a means to initialize the backbone with a starting sequence.

#### Sampling

Given a residue position i and a local environment env_*i*_, around that residue with either just backbone atoms (baseline model) or other residue side-chains as well (conditional model), the sampling procedure is as follows. First, sample a residue type 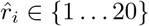 conditioned on the environment. Then, conditioned on the environment and the sampled amino acid type, sample rotamer angle bins 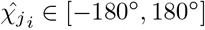 for *j* = 1… 4 in an autoregressive manner. The model in this instance learns a distribution over 24 rotamer bins (7.5° per bin). After a discrete rotamer bin has been sampled, the final rotamer angle 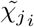, is sampled from a uniform distribution over the bin.

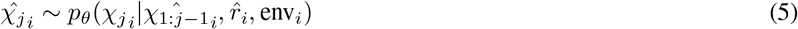

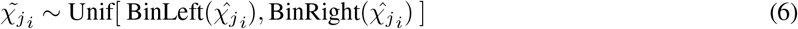

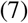

Note that the subsequent autoregressive step conditions on the discrete rotamer bin, not the sampled continuous rotamer angle.

As residues and rotamers are sampled at different positions along the protein backbone, we monitor the negative pseudo-log-likelihood (PLL) of the sequence

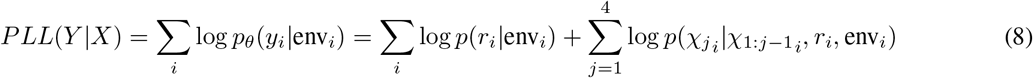

as a heuristic model “energy.” Note that most residues have fewer than four *χ* angles. At sampling time, after a residue type has been sampled, only the corresponding *χ* angles for that residue are sampled. For residues which do not have a particular *χ_j_*, an average value for the log probability of *χ_j_* under the conditional model across the train set and across the rotamer bins is used instead in the PLL calculation.

In this study, we do simulated annealing to optimize the average PLL across residue positions. The acceptance criterion for a step is

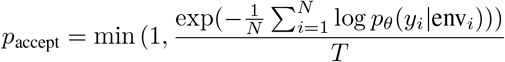

where temperature *T* is annealed over the course of a design run.

In order to speed up convergence, we do blocked sampling of residues. In practice, we draw edges in the graph between nodes where corresponding residues have *C_β_* atoms that are less than 20 Å apart, guaranteeing that non-neighboring nodes correspond to residues that do not appear in each other’s local environments. During sampling, we use greedy graph coloring to generate blocks of independent residues. We then sample over all residues in a block in parallel, repeating the graph coloring every several iterations. Additionally, we restrict the model from designing glycines at non-loop positions, based on DSSP assignment [50, 30].

#### Implementation

The algorithm is implemented in Python, using PyTorch for loading and evaluating the trained networks and PyRosetta for building structural models based on the network predictions for residue type and rotamer angles. Code for the algorithm is available at https://github.com/ProteinDesignLab/protein_seq_des.

#### Runtime

The runtime for our method for sequence design is determined primarily by two steps: (1) sampling residues and rotamer angles and (2) computing the model energy (negative PLL). These times are determined by the speed of the forward pass of the neural network, which is a function of the batch size, the network architecture, and the GPU itself. Note that to compute the model energy, a forward pass of the network is done at each residue position about which the environment has changed.

Annealing for 2500 steps takes between 1 to 2 hours for the native test cases on a computer with 32 GB RAM and on a single GeForce GTX TITAN X GPU, with other processes running on the machine (Dataset S2). *Rosetta-RelaxBB* takes 20-30 minutes per design, while *Rosetta-FixBB* takes 5-15 minutes per design. Compressing the network or modifying the architecture and parallelizing the sampling procedure across more GPUs would improve the overall runtime. In addition, a faster annealing schedule and early stopping of optimization would also reduce the runtime.

### Native test case design experiments

#### Rotamer repacking

Five rounds of rotamer repacking were done on each of the four test case backbones. Repacking is done by fixing the native sequence and randomizing starting rotamers, or using baseline model predictions on backbone atoms only to initialize rotamers. Rotamer prediction at each step and each residue position conditions on the true native residue identity. Model negative PLL averaged by protein length was annealed for 2500 iterations with starting temperature 1 and annealing multiplicative factor 0.995. Commands to reproduce experiments are provided in Dataset S2.

#### Sequence design

Native test case structures *1acf* (2.00 Å), *1bkr* (1.10 Å), *1cc8* (1.022 Å), and *3mx7* (1.76 Å) belong to the beta-lactamase (CATH:3.30.450), T-fimbrin (CATH:1.10.418), alpha-beta plait (CATH:3.30.70), and lipocalin (CATH:2.40.128) topology classes, respectively. We selected these test structures because they span the three major CATH [32, 33] protein structure classes (mostly alpha, alpha-beta, and mostly beta) and because their native sequences were recoverable via structure prediction with Rosetta AbInitio, ensuring they could serve as a positive control for later *in silico* folding experiments. Fifty rounds of sequence and rotamer design were done on each of the four test case backbones. Sequences were initialized via prediction by the baseline model given backbone atoms alone. Model negative PLL averaged by protein length was annealed for 2500 iterations with starting temperature 1 and annealing multiplicative factor 0.995. Commands to reproduce experiments are provided in Dataset S2. Sequences for further characterization were first filtered by specific criteria including all helices capped at the N-terminal, packstat ≥ 0.55 pre and post-RosettaRelax, and for some cases other cutoffs for side-chain and backbone buried unsatisfied hydrogen bond donors or acceptors (Table S1). After filtering, sequences were ranked by model negative PLL for selection. No changes were made to the sequences produced by the model. Top sequences highlighted in Fig. 2I-J are *1acf d3*, *1bkr d2,1cc8 d2,* and *3mx7 d4.*

### TIM-barrel design experiments

The TIM-barrel template backbone is a circularly permuted variant of the reported design sTIM-11 (PDB ID *5bvl*) [40], and the template was prepared using RosettaRemodel with sTIM-11 sequence, circularly permuted and without the originally designed cysteines which did not successfully form a disulfide (C8Q, C181V). sTIM-11 was excluded from the training set is distant in sequence and structure from any known protein, and contains local structural features that differ significantly from naturally occurring TIM-barrels [40]. The original design sTIM-11 was designed using a combination of Rosetta protocols and manual specification of residues. Fifty rounds of sequence and rotamer design were done on the TIM-barrel template backbone. Sequences were initialized via prediction by the baseline model given backbone atoms alone. Model negative PLL averaged by protein length was annealed for 2500 iterations with starting temperature 1 and annealing multiplicative factor 0.995. Residue and rotamer four-fold symmetry was enforced by averaging predicted logits across symmetric positions before normalizing to a discrete distribution and sampling. Commands to reproduce experiments are provided in Dataset S4. Sequences for further characterization were filtered by the following criteria: all helices capped at the N-terminal and packstat ≥ 0.55 pre and post-RosettaRelax. 13 sequences were selected based on structure and sequence metrics (Table S2) and from these 8 sequences were selected based on structural features and a set of simple criteria (Table S3). No changes were made to the sequences produced by the model, except a single cysteine to valine mutation for F1 and F15 in each symmetric subunit (C5V) made ahead of testing for ease of purification, and the valine mutation was ranked highly by the model.

### Design baselines

RosettaDesign [51, 52] is a method for sequence design that has been broadly experimentally validated. [51, 52]. We use Rosetta to design sequences in order to have a point of comparison with the model across the metrics used to evaluate design quality. Further observations about Rosetta performance can be found in the Supplementary Text. The *Rosetta-FixBB* baseline uses the Rosetta packer [51], invoked via the *RosettaRemodel* [53] application, to perform sequence design on fixed backbones. This design protocol performs Monte Carlo optimization of the Rosetta energy function over the space of amino acid types and rotamers [51]. Between each design round, side-chains are repacked, while backbone torsions are kept fixed. Importantly, the Rosetta design protocol samples uniformly over residue identities and rotamers, while our method instead samples from a learned conditional distribution over residue identities. The *Rosetta-RelaxBB* protocol is highly similar to the *Rosetta-FixBB* protocol but performs energy minimization of the template backbone in addition to repacking between design cycles, allowing the template backbone to move. Starting templates for both baselines have all residues mutated to alanine, which helps eliminate early rejection of sampled residues due to clashes. The REF2015 Rosetta energy function was used for all experiments [16, 54]. In order to remove potentially confounding artifacts that emerge during construction and optimization of PDB structures, we additionally relax the test case backbones with constraints to the original atom positions under the Rosetta energy function and run design on these constrained relaxed backbones, as well as for the crystal structure. This step is necessary in order for the Rosetta protocols to in theory be able to recover the native sequence via optimization of the Rosetta energy function.

### Metrics

To assess biochemical properties of interest for the designed sequences we use the following three metrics: (1) packstat, a non-deterministic measure of tight core residue packing [55], (2) exposed hydrophobics, which calculates the solvent– accessible surface area (SASA) of hydrophobic residues [56], and (3) counts of buried unsatisfied backbone (BB) and side-chain (SC) atoms, which are the number of hydrogen bond donor and acceptor atoms on the backbone and side-chains, respectively, that are not supported by a hydrogen bond. We use PyRosetta implementations of these metrics. Backbone relaxes for designs were done with the RosettaRelax application [57, 58, 59, 60], with the ex1 and ex2 options for extra xi and x2 rotamer sampling.

### Structure prediction

#### Rosetta AbInitio

Rosetta AbInitio uses secondary structure probabilities obtained from Psipred [25, 26, 27, 28] to generate a set of candidate backbone fragments at each amino acid position in a protein. These fragments are sampled via the Metropolis-Hastings algorithm to construct realistic candidate structures (decoys) by minimizing Rosetta energy. Native test case runs used Psipred predictions from MSA features after UniRef90 [61] database alignment. TIM-barrel design runs used Psipred predictions directly from sequence. All designs were selected without using any external heuristics, manual filtering, or manual reassignments. We obtained Psipred predictions, picked 200 fragments per residue position [62], and ran 10^4^ trajectories per design. Folding trajectories in Fig. 3, S12 were seeded with native fragments.

#### trRosetta

We also use the trRosetta online server to get structure predictions for designed sequences [22]. trRosetta uses a deep neural network to predict inter-residue distance and orientation distributions from sequence and multiple sequence alignment (MSA) data. These distributions are used to encode restraints to guide Rosetta structure modeling. No homologous structure templates were used for specifying distance constraints in the model-building step. We report alpha-carbon RMSD and GDTMM (Global Distance Test score using Mammoth for structure alignment) as another measure of structure correspondence.

### Protein Purification

Native test case proteins and FXN TIM designs were produced as fusions to an N-terminal 6xHis tag followed by a Tobacco Etch Virus (TEV) cleavage sequence (ENLYFQS). FXC TIM designs were produced as fusions to a C-terminal 6xHis tag. Expression was performed in E. coli BL21(DE3) using the pET24a expression vector under an isopropyl β-D-thiogalactopyranoside (IPTG) inducible promoter. Cultures were induced at OD600 0.5-0.8 by 1mM IPTG at 16°C for 18-20 hours. Proteins were purified by NiNTA-affinity resin (Qiagen). Purity and monomeric state were confirmed using a Superdex 75 increase 10/300 GL column (GE Healthcare) and SDS-PAGE gels. Protein concentration was determined using predicted extinction coefficients and 280 nm absorbance measured on a NanoDrop spectrometer (Thermo Scientific).

### Circular Dichroism Spectroscopy

Circular Dichroism spectra were collected using a Jasco 815 spectropolarimeter with measurements taken in Phosphate Buffered Saline (PBS) using a 1.0mm pathlength cuvette. Wavelength scans were collected and averaged over 3 accumulations. Melting curves were collected monitoring CD signal at 222nm over a range of 25°C to 90°C at 1°C intervals, 1 minute equilibration time and 10 second integration time. For *3mx7* designs, CD signal was monitored at 217nm. Spectra are normalized to mean residue ellipticity (10^3^deg cm^2^ dmol^−1^) from millidegrees. Melting temperatures were determined by fitting a sigmoid function to melting curves.

### Crystallography

#### Screening

F2C and F15C were prepared in a 50mM MES 50mM NaCl pH 6.0 buffer. Crystallization trials were done by screening for crystallization conditions using INDEX (HR2-134), Crystal Screen HT (HR2-130), BCS screen (Molecular Dimensions), MemGold HT-96 (Molecular Dimensions) and a 96-well crystal screen developed at SSRL. A number of crystal hits came from the SSRL screen using Bis Tris (pH 6.0) as the buffer. The best diffracting crystals for F2C were from a well solution consisting of 25% PEG3350, 0.150 M Li2SO4 H2O, and 0.1 M Bis Tris (pH 6.0). The F15C crystals were from a well solution consisting of 30% PEG3350, 0.150 M Mg(OAc)2 4H2O, 0.1 Bis Tris (pH 6.0). The final crystal drops were setup manually as sitting drops in 3 drop crystallization plate that holds 40uL well solution with each drop consisting of 1ul of protein and 1uL of well solution. The F2C crystals appeared after 2 days and the F15C crystals took 3-4 days.

#### Data collection and refinement

Diffraction data was collected at 100° K using Stanford Synchrotron Radiation Lightsource (SSRL) beamlines 12-2 and the Pilatus 6M detector. Data were indexed and integrated using XDS package [63]. Initial phases were obtained by molecular replacement by using the program Phaser [64] and the coordinates of the designed structures as the search model. The high resolution F2C structure was traced with Buccaneer [65] and went through manual fitting using COOT [66] and refinement using REFMAC [67]. The F15C structure refinement involved several cycles of manual model building and refinement. The refinement statistics are provided in Table S5. Structures have been deposited in the Protein Data Bank with accession codes XXX (F2C) and YYY (F15C).

## Contributions

N.A. and P.-S.H. conceived research. N.A. built the algorithm and created the designs with discussions and assistance from R.R.E. and A.D. C.P.P., N.A., and R.R.E conducted experimental validation. I.I.M. solved the protein structures. R.R.E. built the pipelines involving Rosetta. A.D. trained an initial model and contributed to AbInitio experiments with R.R.E. C.P.P., R.R.E, and A.D. contributed to discussion and analysis. P.-S.H. and R.B.A. supervised the research. All authors contributed to the writing and editing of the manuscript.

## Acknowledgements

We thank Tudor Achim for detailed discussion and feedback. We thank Wen Torng, Sergey Ovchinnikov, and Alex E. Chu for helpful discussion and Frank DiMaio for providing a set of decoys from which we selected test case structures. We thank Rohan Kannan for help with running and analyzing rotamer repacking experiments. This project was supported by the U.S. Department of Energy, Office of Science, Office of Advanced Scientific Computing Research, Scientific Discovery through Advanced Computing (SciDAC) program. Additionally, cloud computing credits were provided by Google Cloud. R.B.A. acknowledges support from NIH GM102365, GM61374, and the Chan Zuckerberg Biohub. R.R.E acknowledges support from the Stanford ChEM-H Chemistry/Biology Interface Predoctoral Training Program and the National Institute of General Medical Sciences of the National Institutes of Health under Award Number T32GM120007. A.D. acknowledges support from the National Library of Medicine BD2K training grant LM012409.

Use of the Stanford Synchrotron Radiation Lightsource, SLAC National Accelerator Laboratory, is supported by the U.S. Department of Energy, Office of Science, Office of Basic Energy Sciences under Contract No. DE-AC02-76SF00515. The SSRL Structural Molecular Biology Program is supported by the DOE Office of Biological and Environmental Research, and by the National Institutes of Health, National Institute of General Medical Sciences (P30GM133894). The contents of this publication are solely the responsibility of the authors and do not necessarily represent the official views of NIGMS or NIH.

---

**Movie S1. Examples of model-guided rotamer repacking on native backbone test cases.**

**Movie S2. Examples of model-guided sequence and rotamer design via annealing on native backbone test cases and** *de novo* **TIM-barrel scaffold.**

**SI Dataset S1** (`dataset_Sl.xlsx`)

Training information, including model architecture, excluded ions/ligands, train/test CATH topology classes, and train/test CATH domains

**SI Dataset S2** (`dataset_S2.xlsx`)

Native test case analysis data including rotamer recovery data, sequence design metrics, amino acid distribution data post-design, decoy ranking, runtime raw data, and CD (Circular dichroism) wavelength scans and thermal melt data.

**SI Dataset S3** (`dataset_S3.xlsx`)

TIM-barrel design data, including design metrics and information for designs selected for further characterization.

**SI Dataset S4** (`dataset_S4.zip`)

Other data including design scaffolds, rotamer repacking final models, designed sequence models, positionwise amino acid distributions, CD spectroscopy data, size-exclusion chromatography data, Rosetta AbInitio fragment data, and trRosetta results. Dataset available for download at https://drive.google.eom/file/d/ld8wVjhLzSCKfwmHeVoeuYV_JQXkPXysN/view?usp=sharing

Code for running sequence design algorithm available at https://github.eom/ProteinDesignLab/protein_seq_des.

## Supplementary text

### Classifier performance

For the conditional residue prediction task, our classifier gives an improvement of 14.7% over [34] (42.5%), and similar performance to [37] (52.4%), [35] (56.4%) and [36] (58.0%). We note that we do not use the same train/test sets as these studies. Another study uses a similar 3D CNN model for ΔΔG prediction [68]. The baseline model has a 33.5% test set accuracy. The model often confuses large hydrophobic residues phenylalanine (F), tyrosine (Y), and tryptophan (W), and within this group, more often confuses Y for F which are similarly sized compared to the bulkier W. Looking at residue-specific accuracies (Fig. S1C, top), on the whole, the conditional model is more certain about hydrophobic/non-polar residue prediction compared to hydrophilic/polar. Both the conditional and baseline models do especially well at predicting glycine and proline residues, both of which are associated with distinct backbone torsion distributions (Fig. S1C).

The conditional model is also trained to predict binned rotamer torsion (χ) angles in an autoregressive fashion. Across 24 bins, the test set accuracy of the model to within less than 15 degrees is 52.7%, 43.6%, 26.8%, and 30.6% for *χ*_1_-*χ*_4_, respectively (Fig. S1D). Unlike rotamer scoring methods [39, 69], where the network is trained to score contextually correct rotamers from others, this model is simply trained to predict all of the rotamer angles, conditioned on context.

### Generalization to unseen topologies – design baselines

We assess the performance of our sequence design method by comparison with two Rosetta baselines. The first, *Rosetta-FixBB*, is a fixed-backbone Rosetta design protocol, and the second, *Rosetta-RelaxBB*, interleaves backbone relaxes between fixed-backbone design cycles, allowing the template backbone to move. The *Rosetta-FixBB* protocol provides the most relevant comparison to our method as both operate on fixed backbones. However, *Rosetta-RelaxBB* is the more commonly used mode of sequence design, generating solutions that are more variable than those of *Rosetta-FixBB.* We have therefore included designs from both methods for comparison.

On average, Rosetta is more sensitive to the backbone than the model for rotamer repacking: for the test case *1acf*, Rosetta does well once the crystal structure backbone has been relaxed under the Rosetta energy function with the native sequence and rotamers in place; our model, however, is robust to the deviations in the backbone and performs similarly in both cases (Fig. S3A).

The model recapitulates the native sequence to a similar extent as *Rosetta-FixBB* and to a better extent than *Rosetta-RelBB* (Fig. 2C and S3B). While the *Rosetta-FixBB* designs are convergent, the model designs are more variable in terms of inter-sequence overlaps across designs (Fig. 2D and S4B). However, both the model designs and Rosetta designs converge to similar core (low solvent-accessible surface area) regions, indicating that perhaps the need for a well-packed core in combination with volume constraints determined by the backbone topology might limit the number of feasible core designs.

Model-designed sequences adhere to an amino acid distribution similar to the native sequence and its aligned homologs (Fig. S4A), with the most pronounced difference being an apparent overuse of lysine (K) and glutamate (E) and under-use of rarer residues (H, N, Q, W, C) relative to homologous sequences; *Rosetta-FixBB* designs also overuse K and E and under-use H, W, N, and C relative to the native MSA distribution. Like the model, Rosetta protocols match the native distribution well, with a tendency to overuse small hydrophobic residues, in particular alanine (A), valine(V), and isoleucine (I).

Sequences designed onto the test case backbones should ideally have the following key biochemical properties [70, 71]: (1) core regions should be tightly packed with hydrophobic residues, a feature that is important for driving protein folding and maintaining fold stability [72, 73, 74]; (2) designs should have few exposed hydrophobic residues, as these incur an entropic penalty as a result of solvent ordering under the hydrophobic effect and an enthalpic penalty from disrupting hydrogen bonding between polar solvent molecules [75, 76, 56], making it energetically favorable to place polar residues at solvent-accessible positions and apolar residues at buried positions; (3) If polar residues are designed in core regions, they should be supported by hydrogen bonding networks [77, 9].

We compare our designs to the *Rosetta-FixBB* and *Rosetta-RelaxBB* protocol designs, as well as the native idealized structure, across key metrics (Fig. S5). Since the designs have less than 50% sequence overlap with the native on average, we also include data for the native sequence with either 50% or 75% of the residues randomly mutated; these perturbed sequences serve as negative controls and indicate expected performance for likely non-optimal deviations from the native sequence.

We also report performance across these metrics for designed sequences relaxed with the *RosettaRelax* protocol. This procedure allows the backbone and rotamers to move to adopt a lower Rosetta energy conformation. Relaxing the designs allows us to assess whether alternate rotamers and deviations of the starting protein backbone can lead to an improvement in performance across the metrics considered.

1. *Packstat.* Model designs tend to have a lower packstat score compared to to the native sequence and the Rosetta designs (Fig. S5A); however, on average the packstat scores are still higher relative to random perturbations of the native sequence. While this trend seems to indicate that the model designs are less optimal than the Rosetta designs, when we look at the model designs post-relax (Fig. S5B), the packstat scores improve and better match those of the native sequence and Rosetta designs, while the perturbed sequence packstat scores remain low. At the same time, the post-relax alpha-carbon backbone RMSDs between design methods are also comparable (Fig. S6E). These results suggest that the designed sequences do tightly pack core regions, as slight movements of the backbone and repacked rotamers for model designs give well-packed structures.
2. *Exposed hydrophobics.* Model designs in general do not place hydrophobic residues in solvent-exposed positions, similar to the Rosetta designs (Fig. S5C). This trend in the model designs is largely due to the relative abundance of cytosolic proteins in the PDB compared to membrane proteins, which would in general have hydrophobic regions exposed to the membrane lipid environment. The native sequence for test case *3mx7* has many exposed hydrophobic residues, suggesting that the native protein might bind to a target, forming a hydrophobic interface.
3. *Buried unsatisfied backbone atoms.* For all of the test cases except *1bkr,* model designs have similar or fewer unsatisfied backbone polar atoms compared to the native sequence (Fig. S5D-E). For *1bkr,* although the average number of unsatisfied backbone polar atoms is greater than that of the native sequence or Rosetta designs, the distribution is fairly wide, indicating that there are many designs that have fewer unsatisfied backbone polar atoms compared to the native sequence. However, some of the 50% perturbed sequences have fewer unsatisfied backbone polar atoms than the native, suggesting that this metric alone is not sufficient for selecting or rejecting designs.
4. *Buried unsatisfied side-chain atoms.* Model designs across test cases have few unsatisfied buried side-chain polar atoms, similar to the native sequences and Rosetta designs (Fig. S5F,G). This indicates that over the course of sampling, side-chains that are placed in buried regions are adequately supported by backbone hydrogen bonds or by design of other side-chains that support the buried residue.

We use Rosetta AbInitio [51] and trRosetta [22] structure prediction to assess the recoverability of native test case backbones. Selected model designs and the top Rosetta-FixBB design (best of 50 designs ranked by Rosetta energy) are recoverable by both methods, as are some of the top Rosetta-RelBB designs (best of 50 designs ranked by Rosetta energy) (Fig. S7, S8). The 50% perturbed negative control (best of 50 designs ranked by Rosetta energy) sequences are not recoverable under Rosetta, but seem to be recoverable under trRosetta. This is likely due to the fact that native homologs can be found via multiple sequence alignment MSA) despite the sequence perturbation, and the distance and orientation distribution prediction network, having been trained on native structures, can extrapolate from the sequence coevolution signal that is retained. Still, the model designs are better recovered in terms of alpha-carbon RMSD (Å) and GDTMM score than the 50% perturbed sequence.

#### Heuristic structure energy

Although the Rosetta energy is not optimized during our design procedure, many of the designs have low Rosetta energy relative to perturbed sequence baselines (Fig. S6A). After relaxing the model designs, their Rosetta energy decreases to match the native backbone, with relatively low subsequent backbone deviation; in contrast, sequences that are 50% similar to native, with random perturbations, have a high Rosetta energy, even after relax (Fig. S6B) The model energy too is similarly low for the model designs and the Rosetta designs, indicating that the model energy as a heuristic roughly matches Rosetta in differentiating low-energy vs. high-energy designs (Fig. S6D). Overall, the alpha-carbon RMSD distributions post-RosettaRelax for the model designs correspond with the native backbone relaxed distributions (Fig. S6E).

**Figure S1:**
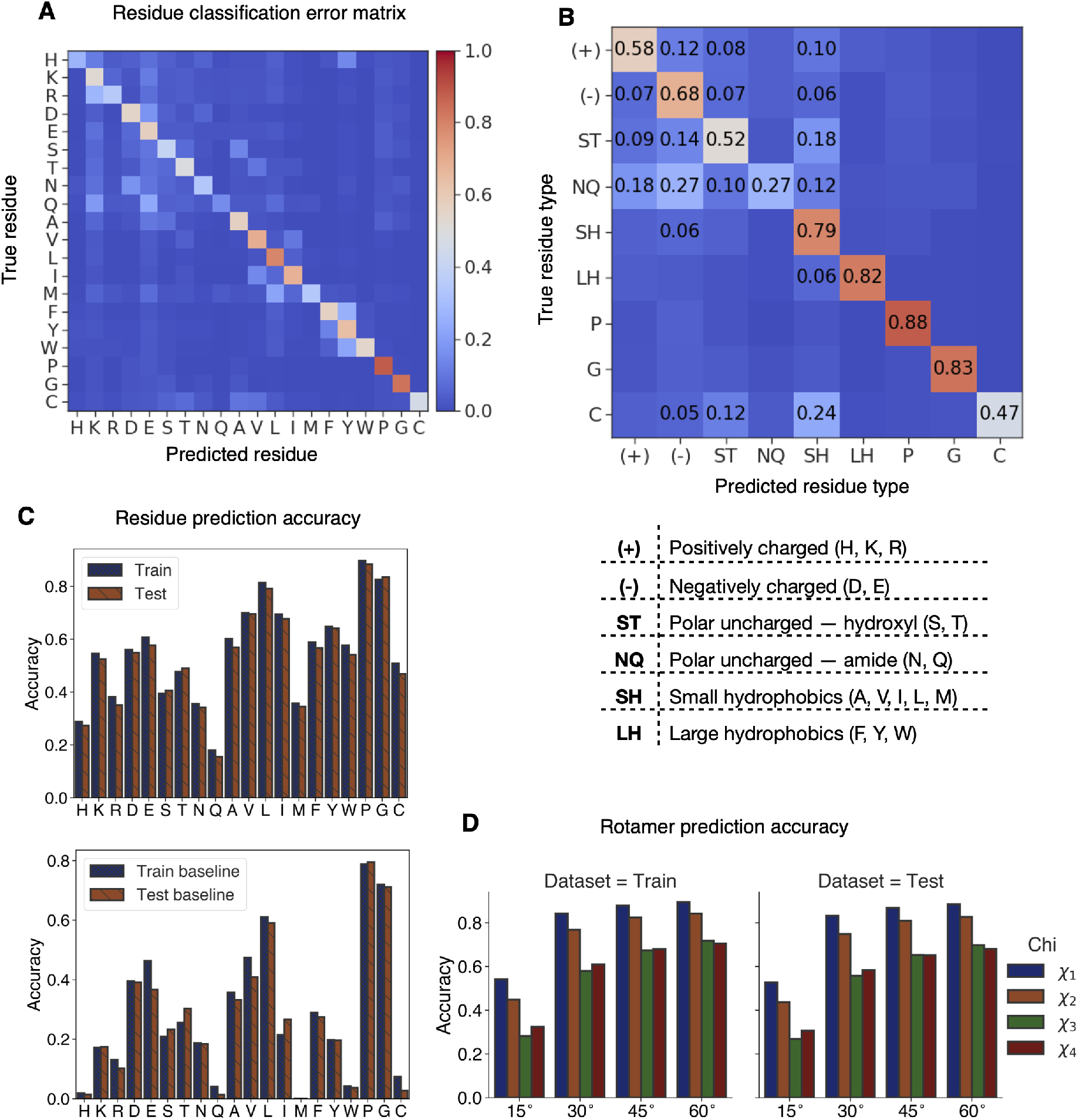
Classifier performance and learned rotamer distributions. **(A)** Classifier error matrix for individual residue prediction on test set data and for **(B)** prediction of groups of biochemically similar residues. **(C)** Residue-specific prediction accuracy for train and test set data for the (top) conditional model and (bottom) baseline model. **(D)** Rotamer prediction accuracy to within 15, 30, 45, and 60 degrees across the train and test set.

**Figure S2:**
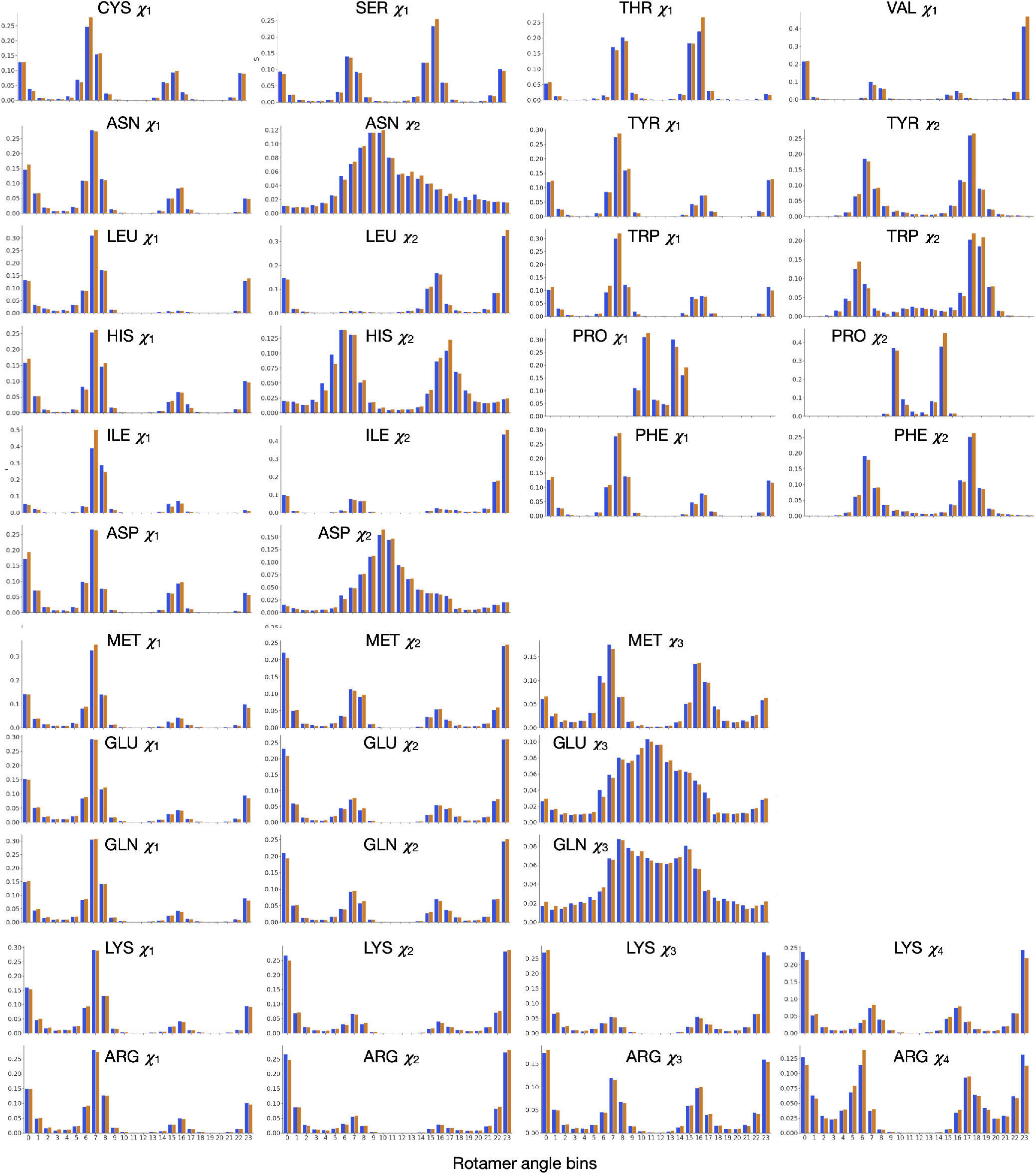
Learned residue-specific rotamer distributions. Binned rotamer distributions for the native test set domains (blue) versus the model predicted distributions (orange). Native distributions are the normalized empirical rotamer distributions for each residue across the test set. Model distributions are the average of the network-predicted distributions across the test set examples.

**Figure S3:**
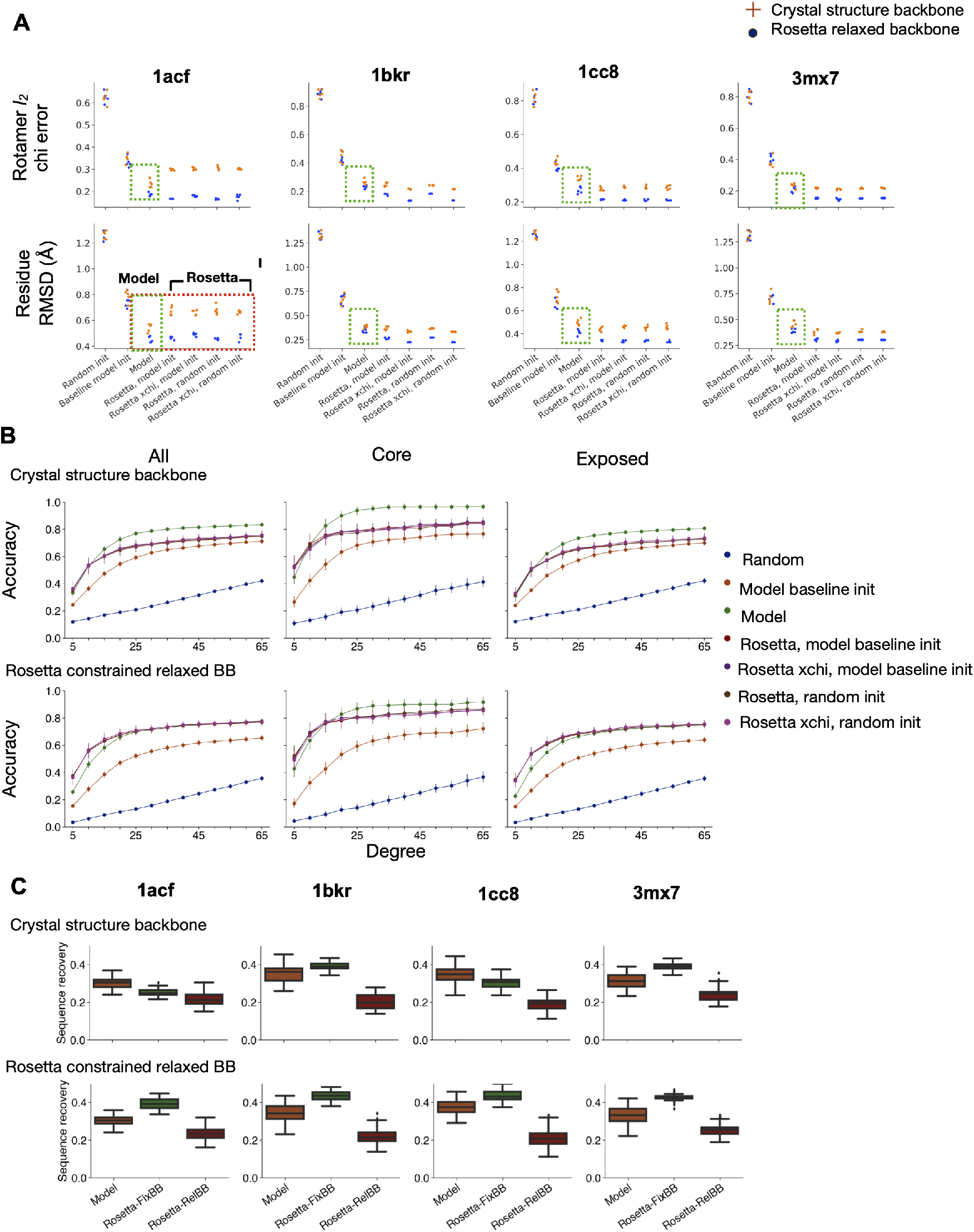
Rotamer and native sequence recovery. (**A – B**) Rotamer recovery. *n* = 5 for all methods for each test case. Data shown for rotamer recovery on crystal structure backbone, as well as Rosetta constrained relaxed backbones. Random init – random initialization of chi angles. Baseline model init – initialization with rotamer prediction by baseline classifier model from backbone alone. Model – rotamer sampling and annealing with conditional model. Rosetta protocols were run either with or without extra *χ*_1_ and *χ*_2_ sampling (xchi). **(A)** (Top) Average *l*_2_ chi angle error across design methods. (Bottom) Average residue side-chain RMSD (Å). **(B)** Rotamer recovery accuracy as a function of degree cutoff for all residues, core residues, and solvent-exposed residues (*n* = 20). **(C)** Native sequence recovery across test case crystal structures (*n* = 50, each). (Top) Design on crystal structures. (Bottom) Design on Rosetta constrained relaxed backbones.

**Figure S4:**
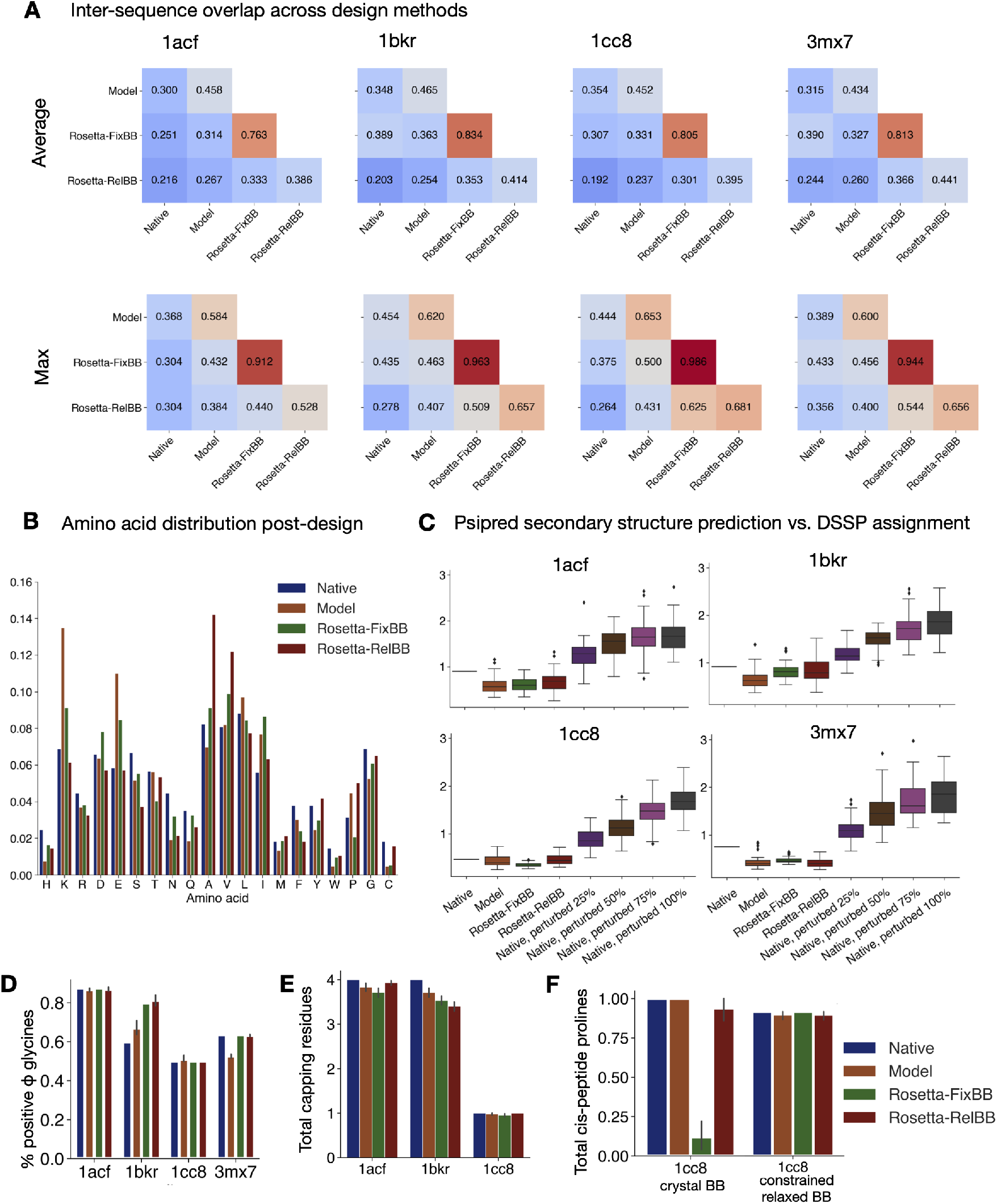
Assessing model designs in comparison to Rosetta baselines. **(A)** Inter-sequence overlap. Average (top) and maximum (bottom) inter-sequence fractional overlap within and across top sequences from each design method. **(B)** Amino acid distribution post-design relative to native sequence and aligned homologs (max. 500 hits per sequence). MSAs obtained using PSI-BLAST v2.9, the NR50 database, BLOSUM62 with an E-value cutoff of 0.1. **(C)** Psipred secondary structure prediction for designed sequences. Cross-entropy of Psipred prediction (helix, loop, sheet) from sequence alone with respect to DSSP assignments [25, 26, 27, 28, 29, 30]. **(D)** Percent glycines at positive *φ* backbone positions across test cases (*n* = 50, each). **(E)** Capping residue placement. Average number of N-terminal helical capping residues across designs for test cases with capping positions (*n* = 50, each). **(F)** Total number of cis-peptide prolines (*ω*_*i*-1_ < 15) for *1cc8* for designs on crystal structure backbone vs. Rosetta constrained relaxed backbone (*n* = 50).

**Figure S5:**
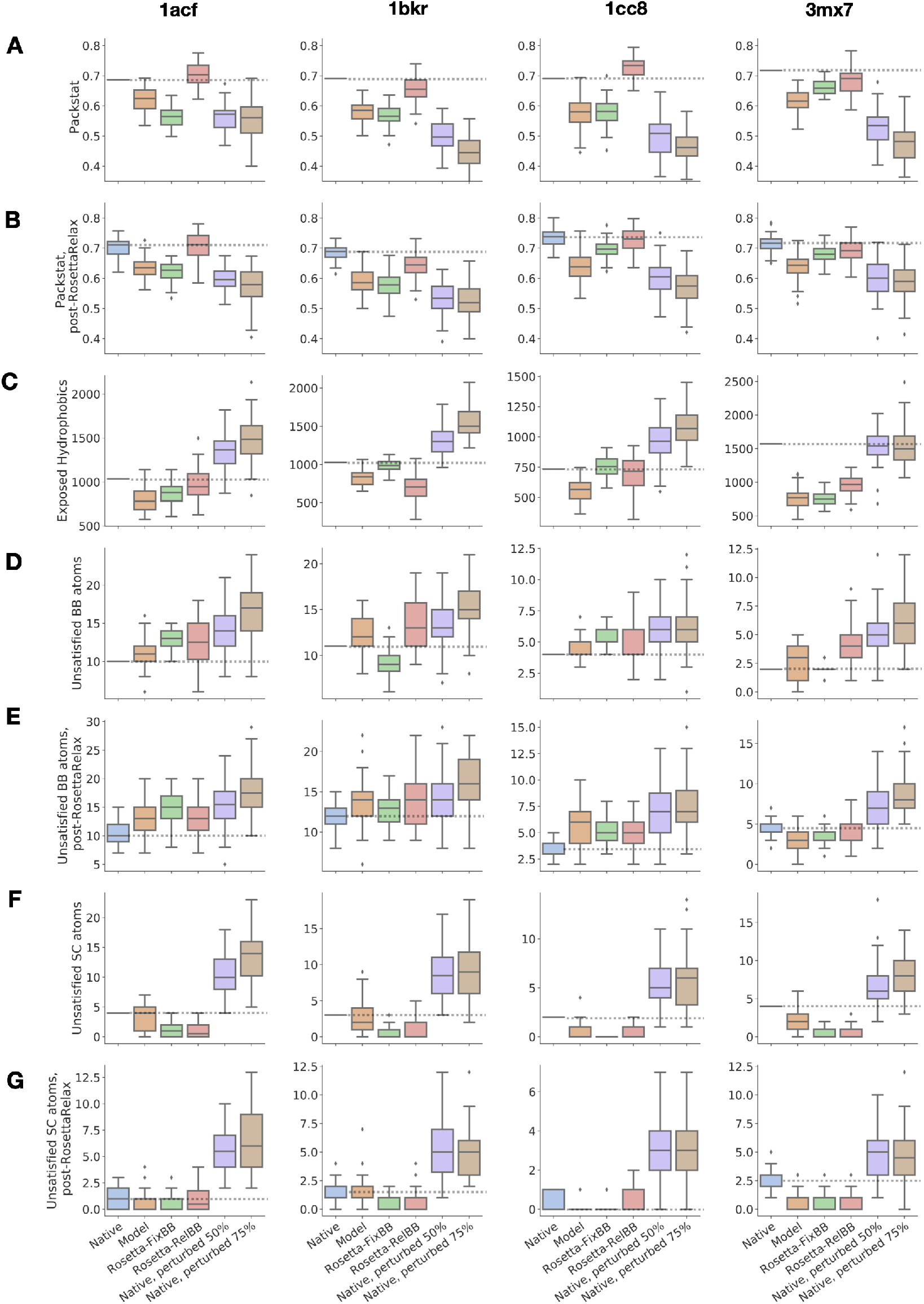
Biochemical metrics of interest for designs on crystal structures. Designs (*n* = 50) compared to native crystal structure (*n* = 1) or distribution of relaxed native structures (*n* = 50). 50% and 75% mutated native sequences included as negative controls. (**A-B**) Packstat, a measure of core residue packing [55] pre- **(A)** and post- **(B)** RosettaRelax [57, 58, 59, 60]. **(C)** Total solvent-accessible surface area (SASA) of exposed hydrophobic residues [56]. (**D-E**) Number of buried unsatisfied polar backbone (BB) atoms pre- **(D)** and post- **(E)** relax. (**F-G**) Number of buried unsatisfied polar side-chain (SC) atoms pre- **(F)** and post- **(G)** relax.

**Figure S6:**
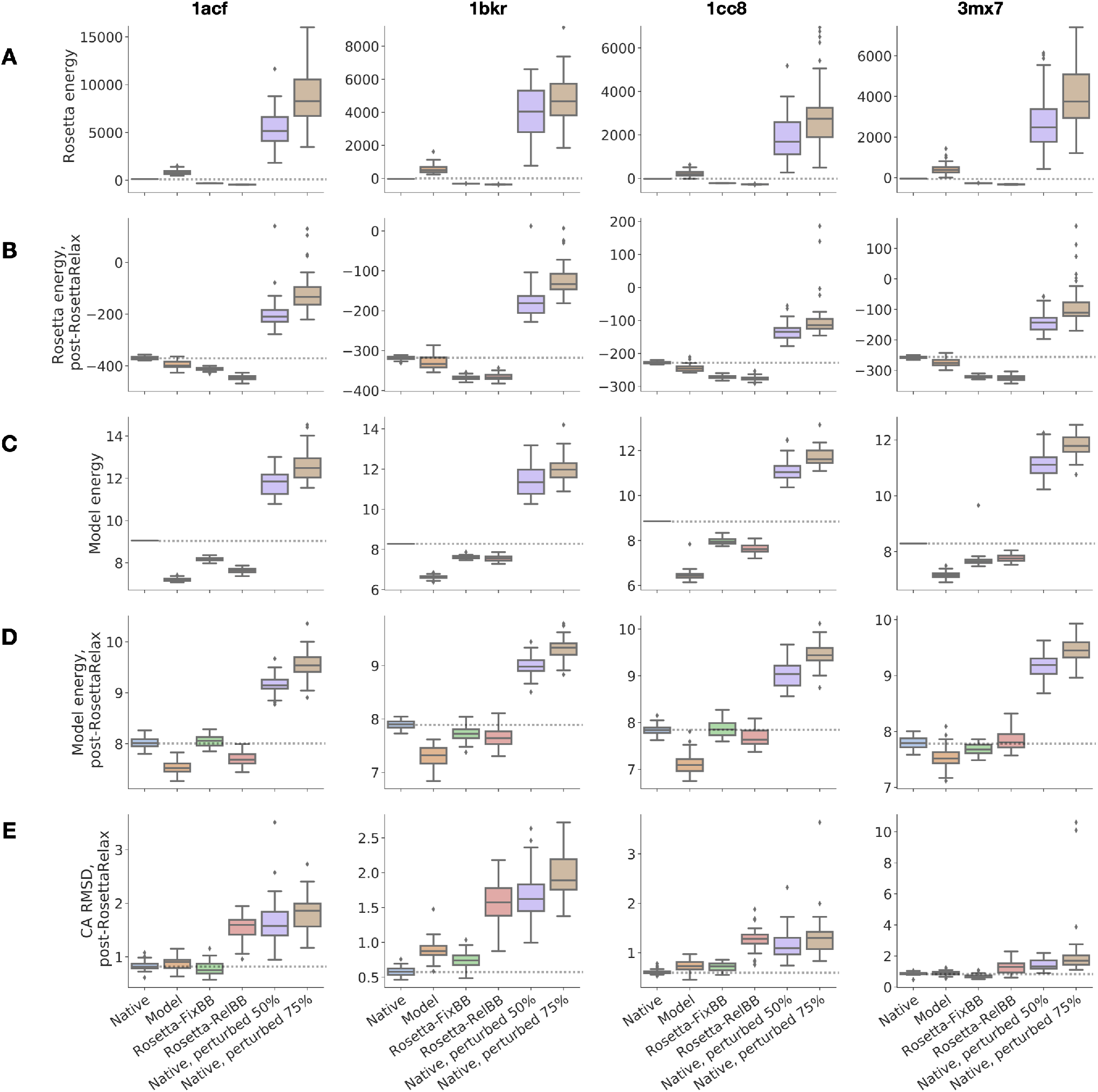
Energy under the model and under the Rosetta energy function pre- and post-RosettaRelax. 50% and 75% mutated native sequences included as negative controls. Designs (*n* = 50) compared to native crystal structure (*n* = 1) or distribution of relaxed native structures (*n* = 50). (**A-B**) Rosetta energy pre- **(A)** and post- **(B)** RosettaRelax. (**C-D**) Model energy (negative pseudo-log-likelihood, normalized by protein length) pre- **(C)** and post- **(D)** relax. **(E)** Alpha-carbon RMSD (Å) post-RosettaRelax.

**Figure S7:**
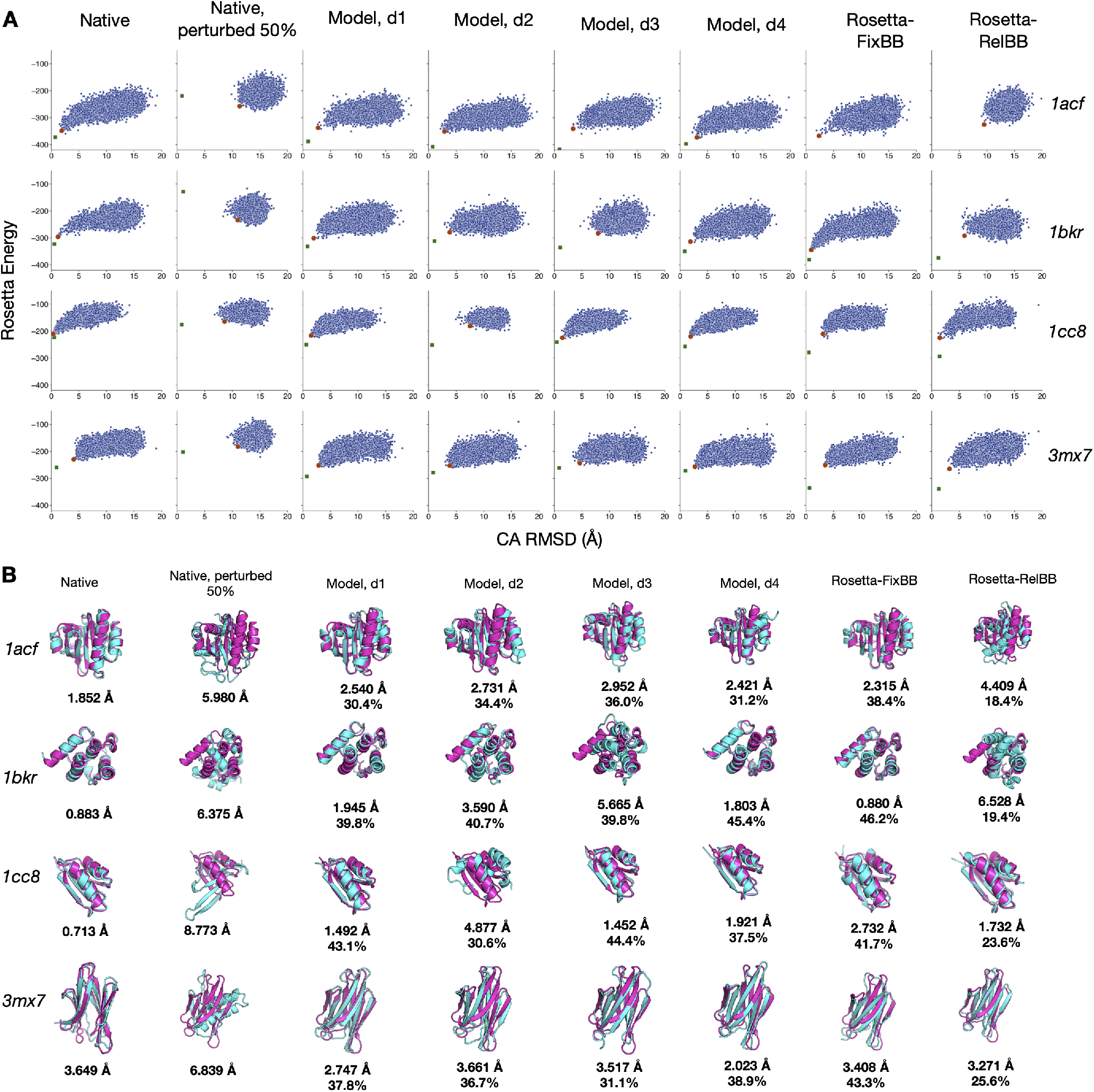
Rosetta AbInitio structure prediction of designed native test case sequences and Rosetta-FixBB and Rosetta-RelBB baselines (best of 50 sequences by Rosetta energy). 50% randomly perturbed native sequences included as a negative control (best of 50 sequences by Rosetta energy). **(A)** Rosetta energy vs. RMSD (Å) to native funnel plots. Selected structure with best summed rank of template-RMSD and Rosetta energy is shown in orange. The designed sequence after RosettaRelax is shown in green. **(B)** Folded structure with best summed rank of template-RMSD and Rosetta energy across 10^4^ folding trajectories. Decoys (blue) are aligned to the native backbones (pink). Sequence identity and RMSD (Å) compared to native are reported below the structures.

**Figure S8:**
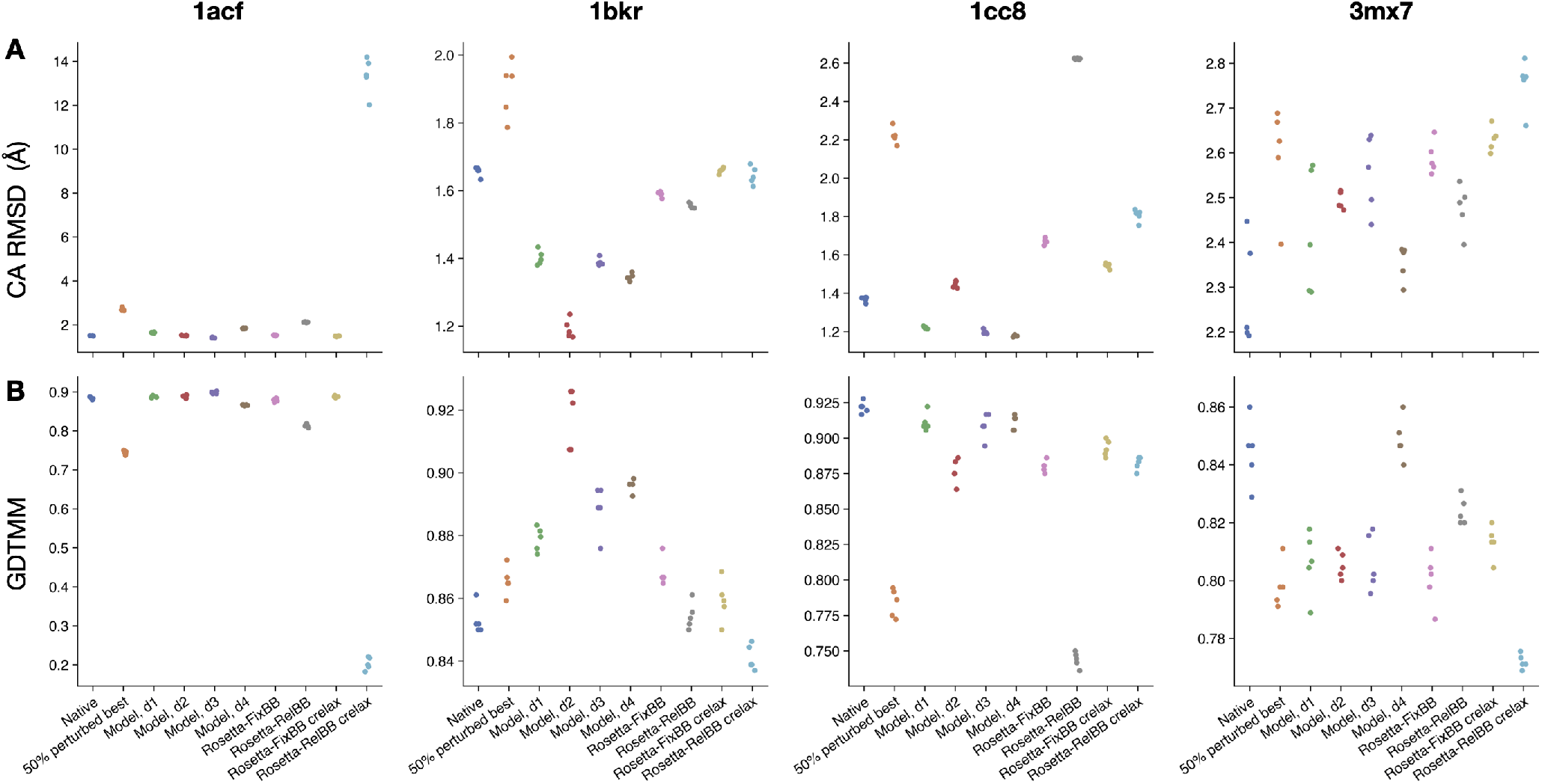
trRosetta structure prediction of designed native test case sequences and baselines. Data for 5 predicted 3D structure models. Data shown for native sequence, 50% perturbed native sequence (best of 50 sequences in Rosetta energy), model designs, best of 50 Rosetta-FixBB and Rosetta-RelBB designs on crystal structure (in terms of Rosetta energy), and best of 50 Rosetta-FixBB and Rosetta-RelBB designs on the constrained relaxed crystal backbone. **(A)** Alpha-carbon (CA) RMSD (Å) to native. **(B)** GDTMM score (higher is better).

**Figure S9:**
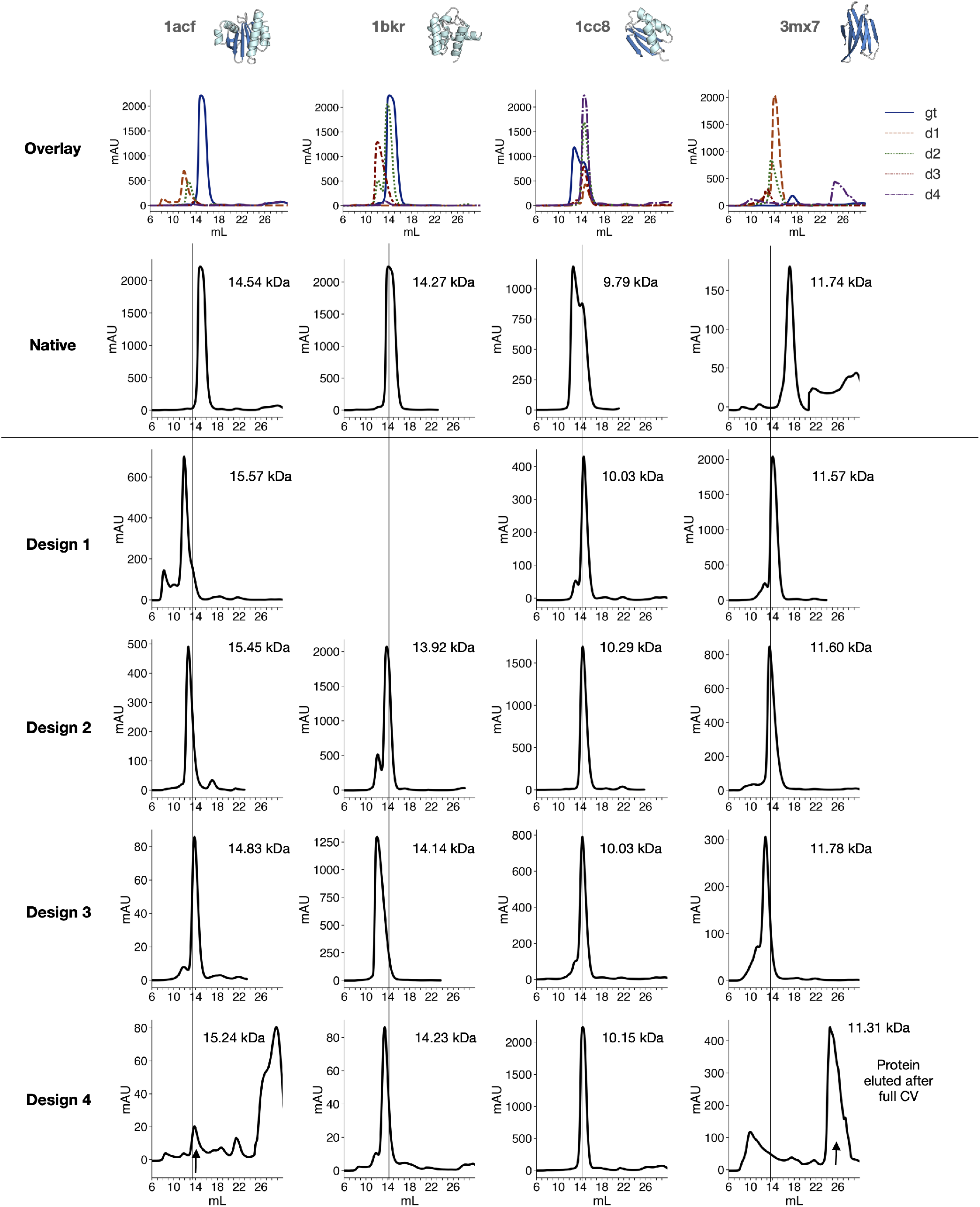
Determining purity and monomeric state for native test case designs via size-exclusion chromatography (Superdex 75). Vertical line shows expected elution volume based on molecular weight of the native sequence. Some proteins seem to interact with the column, eluting later than expected *(3mx7* native, *3mx7* d4). No data for *1bkr* d1 due to low protein expression. Right-most peak for *1acf* d4 is imidazole.

**Figure S10:**
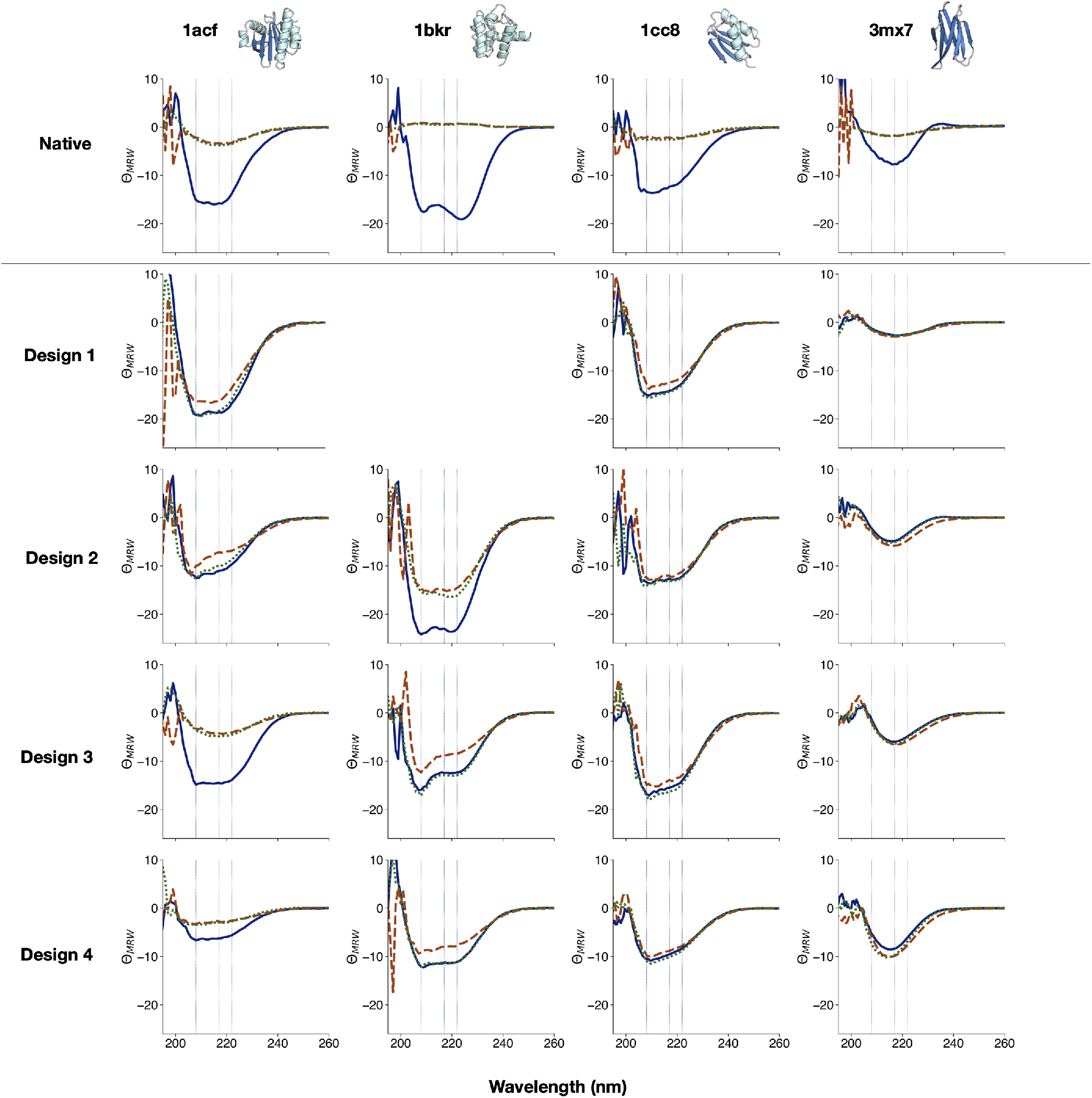
Circular dichroism (CD) spectroscopy wavelength scans for select model designs on native test case backbones. Mean residue ellipticity *Θ_MRW_* (10^3^deg cm^2^ dmol^−1^) for CD wavelength scans at 20°C (blue, solid), melted at 95°C (orange, dashed), and cooled again to 20°C (green, dashed). No data for *1bkr* d1 due to low protein expression. Top sequences highlighted in main paper in Fig. 2I-J are *1acfd3,1bkr d2,1cc8 d2,* and *3mx7 d4.*

**Figure S11:**
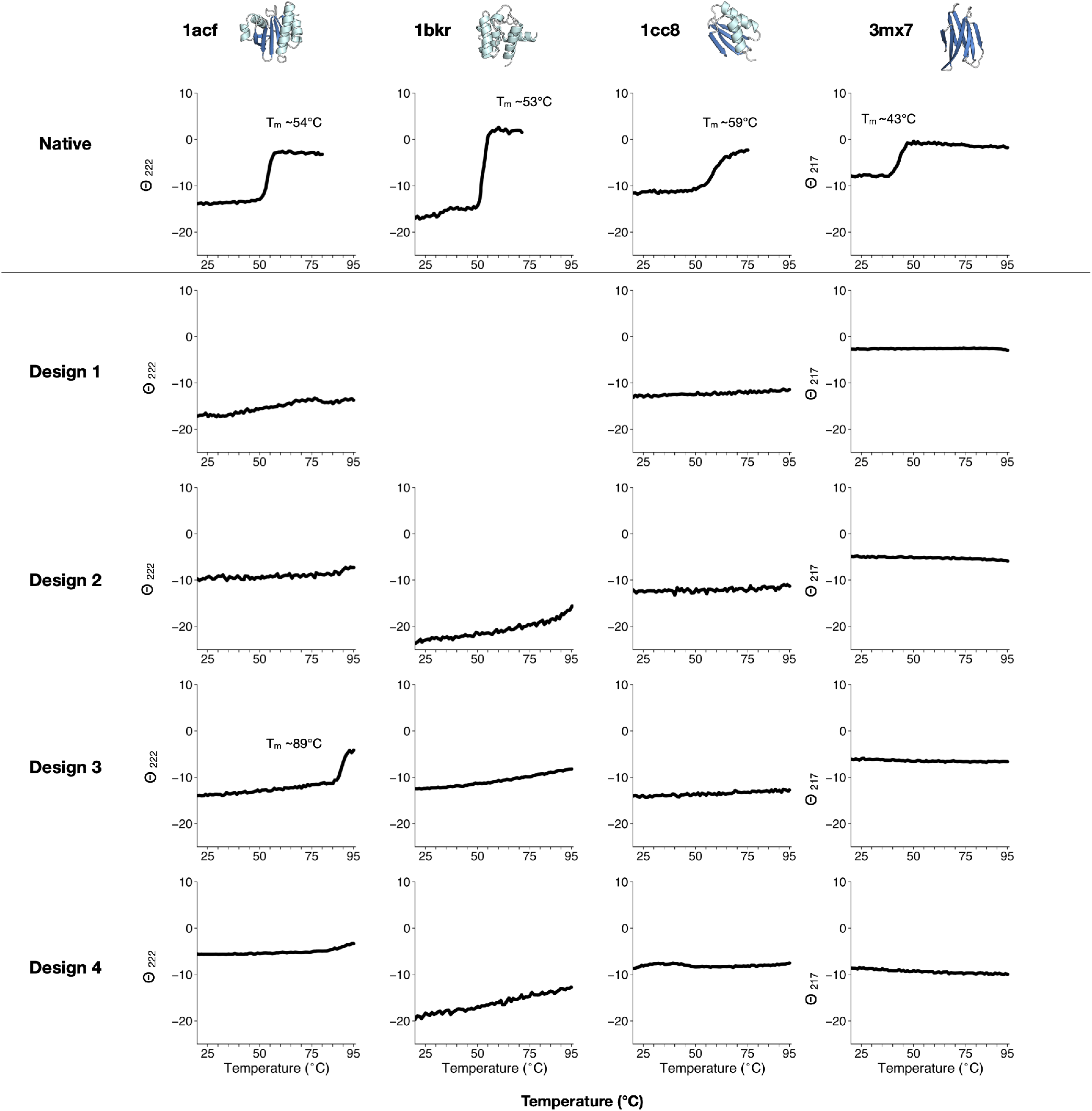
Thermal melting curves for native test case designs monitoring circular dichroism (CD) signal *Θ_MRW_* (10^3^deg cm^2^ dmol^−1^) at 222nm or 217nm for *3mx7* designs. No data for *1bkr* d1 due to low protein expression. For melts with a cooperative transition, melting temperatures *T_m_* listed. Top sequences highlighted in main paper in Fig. 2I-J are *1acfd3,1bkr d2,1cc8 d2,* and *3mx7 d4.*

**Figure S12:**
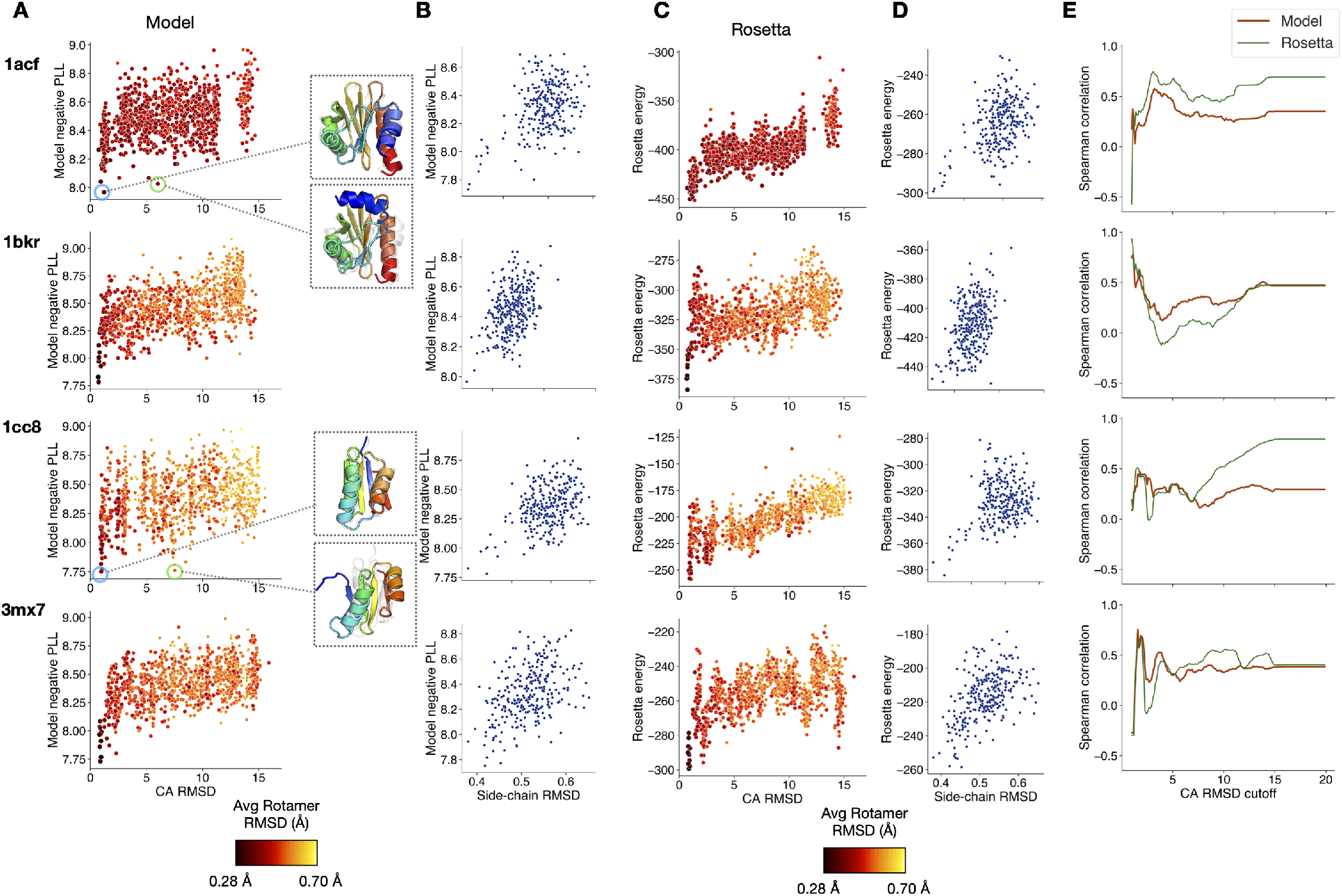
Decoy ranking for test cases. **(A)** Model energy (negative PLL) vs. alpha-carbon RMSD (Å) for Rosetta AbInitio folded structures (decoys). Points are colored by average side-chain RMSD (Å) to native. (Inset) Select structures rendered to visualize alternative minima under model ranking for *1acf* and *1cc8:* for *1acf* an alternative N-terminal helix conformation and for *1cc8* an alternative pattern of beta strand pairing. **(B)** Model negative PLL of low backbone RMS structures (CA RMSD < 5 Å) vs. average side-chain RMSD (Å). **(C)** Rosetta energy vs. alpha carbon RMSD (Å) for folded structures. Points are colored by average side-chain RMSD. **(D)** Rosetta energy of low backbone RMS structures (CA RMSD < 5 Å) vs. average side-chain RMSD (Å). **(E)** Spearman rank correlation between model negative PLL/ Rosetta energy and structure alpha-carbon RMSD (Å) as a function of increasing RMSD cutoff. In the low RMS regime (< 5-10 Å), the model and Rosetta are able to rank low RMS structures to a similar extent.

**Figure S13:**
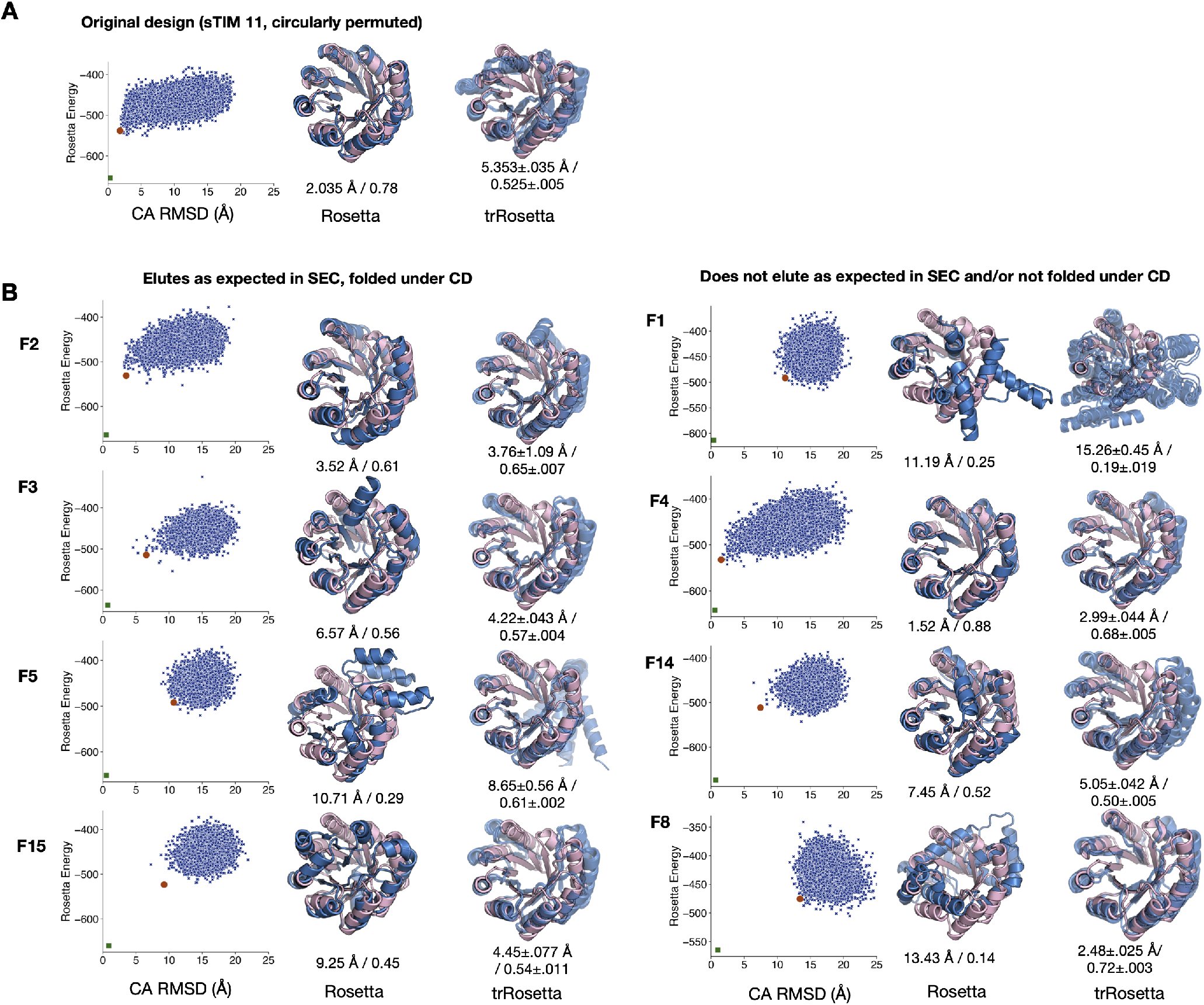
Structure prediction for **(A)** original sTIM-11 sequence (circularly permuted with mutations C8Q, C181V) and (**B**) TIM-barrel model designs. **(Left)** Rosetta AbInitio structure prediction Rosetta energy vs. RMSD (Å) to native funnel plots. Selected structure with best summed rank of template-RMSD and Rosetta energy is shown in orange. The design after RosettaRelax is shown in green. **(Center)** Folded structure with best summed rank of template-RMSD and Rosetta energy across 10^4^ folding trajectories. Decoys (blue) are aligned to the target backbone (pink). Backbone alpha-carbon RMSD (Å) and GDTMM are reported below the structures. **(Right)** trRosetta prediction results. 5 models (blue) overlaid on target scaffold (pink). Backbone alpha-carbon RMSD (Å) and GDTMM below structures.

**Figure S14:**
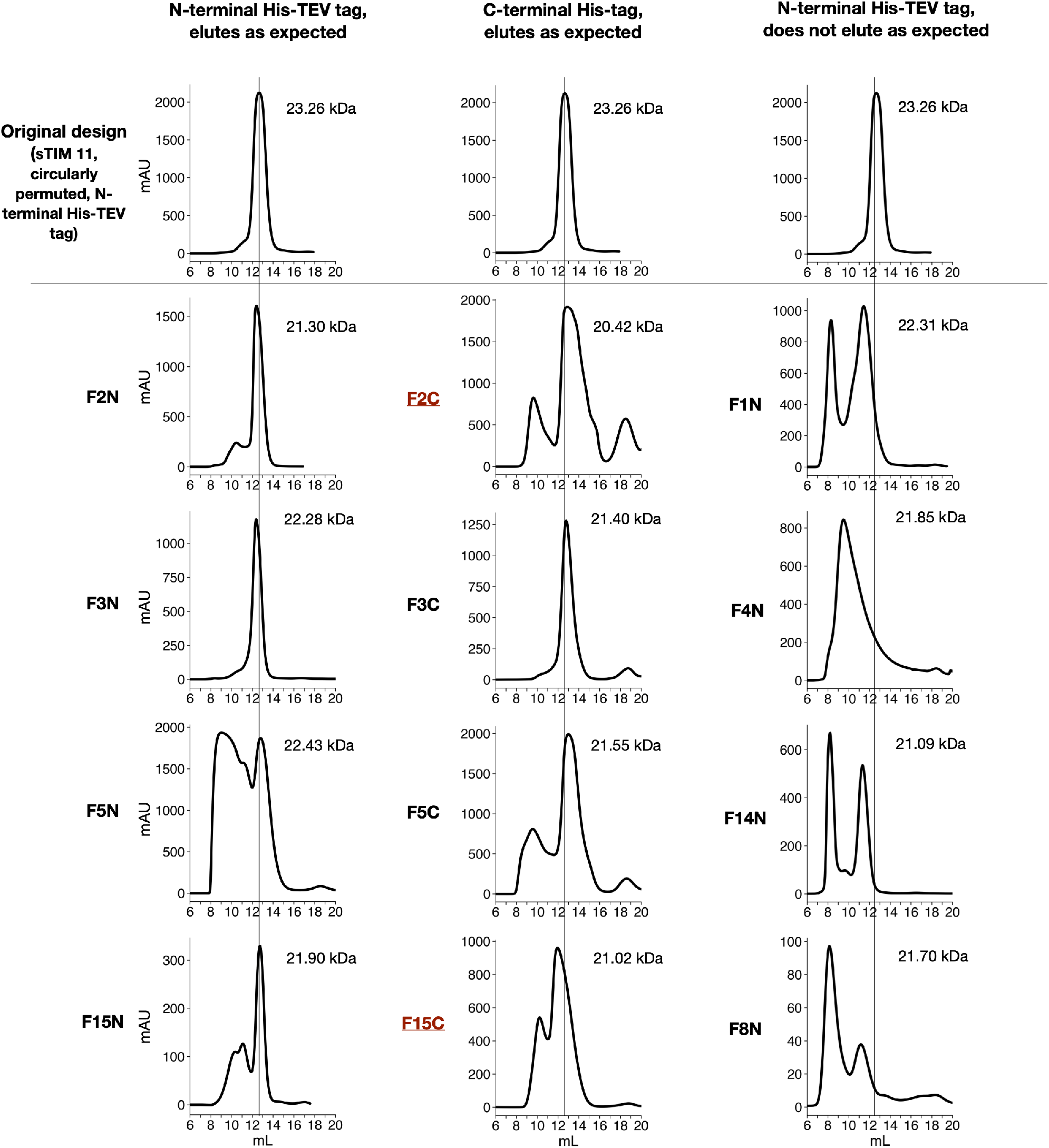
Determining purity and monomeric state for model TIM-barrel designs designs via size-exclusion chromatography (Superdex 75). Crystal structure constructs F2C and F15C highlighted in red. Vertical line shows expected elution volume based on molecular weight of the sTIM-11 sequence. **(Left)** FXN N-terminal His-TEV tagged constructs that eluted as expected. **(Center)** FXN C-terminal His-tagged constructs. **(Right)** FXN N-terminal His-TEV tagged constructs that did not elute as expected.

**Figure S15:**
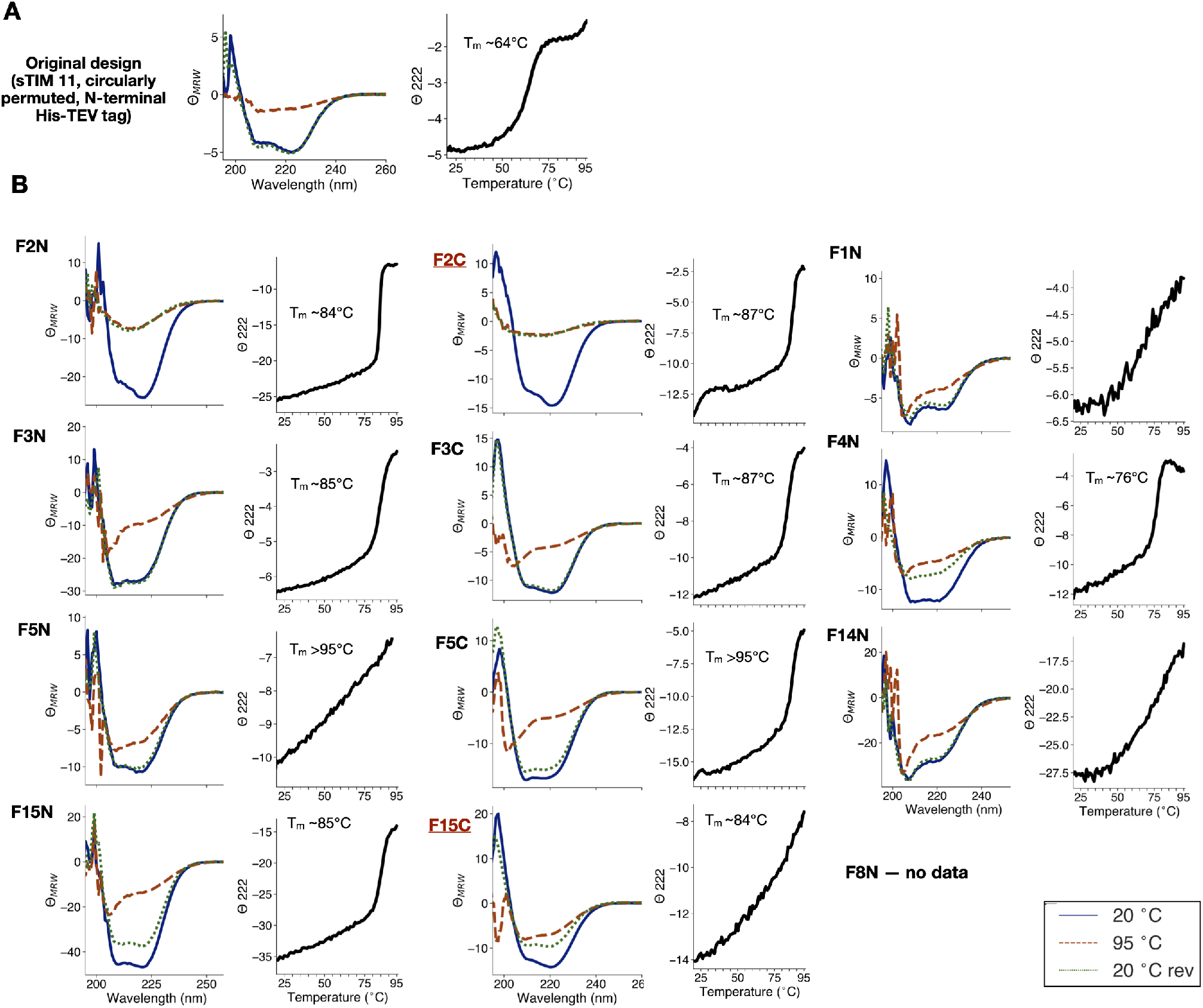
(**A – B**) Circular dichroism (CD) data for (**A**) original sTIM-11 sequence, (circularly permuted with mutations C8Q, C181V) with N-terminal His-TEV tag and (**B**) TIM-barrel model designs: FXN (N-terminal His-TEV tag), FXC (C-terminal His tag). Crystal structure constructs F2C and F15C highlighted in red. **(Left)** Mean residue ellipticity Θ_MRW_ (10^3^deg cm^2^ dmol^−1^) for CD wavelength scans at 20°C (blue, solid), melted at 95°C (orange, dashed), and cooled again to 20°C (green, dashed). **(Right)** Thermal melting curves monitoring CD signal *Θ_MRW_* (10^3^deg cm^2^ dmol^−1^) at 222nm.

**Figure S16:**
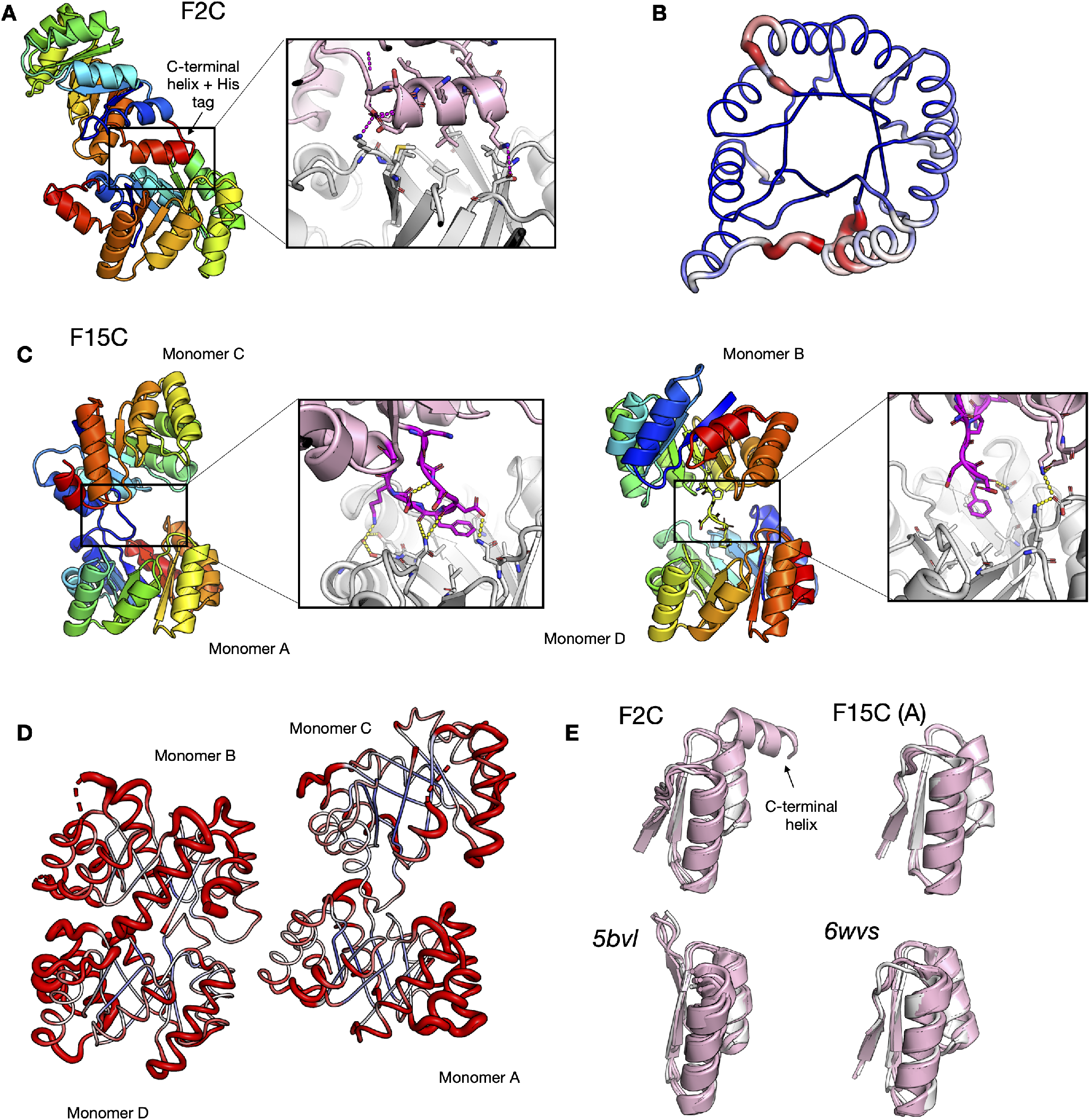
Additional data on TIM-barrel crystal structures. **(A)** The C-terminal helix of the TIM-barrel dislodges from the barrel and makes contact with the bottom of the barrel of an adjacent protein. (Inset) The C-terminal helix plugs into hydrophobic residues in the barrel, while also making polar contacts. **(B)** F2C crystal structure colored by B-factor. **(C)** For two monomers of the tetrameric asymmetric unit of F15C (Monomers C, B), the *β-α* loop to the long helix for one subunit dislodges to interact with the top of the barrel of an adjacent monomer (Monomers A, D) **(D)** Tetrameric asymmetric unit for F15C crystal, colored by B-factor. **(E)** Symmetric units (pink) overlaid with design template (gray).

**Figure S17:**
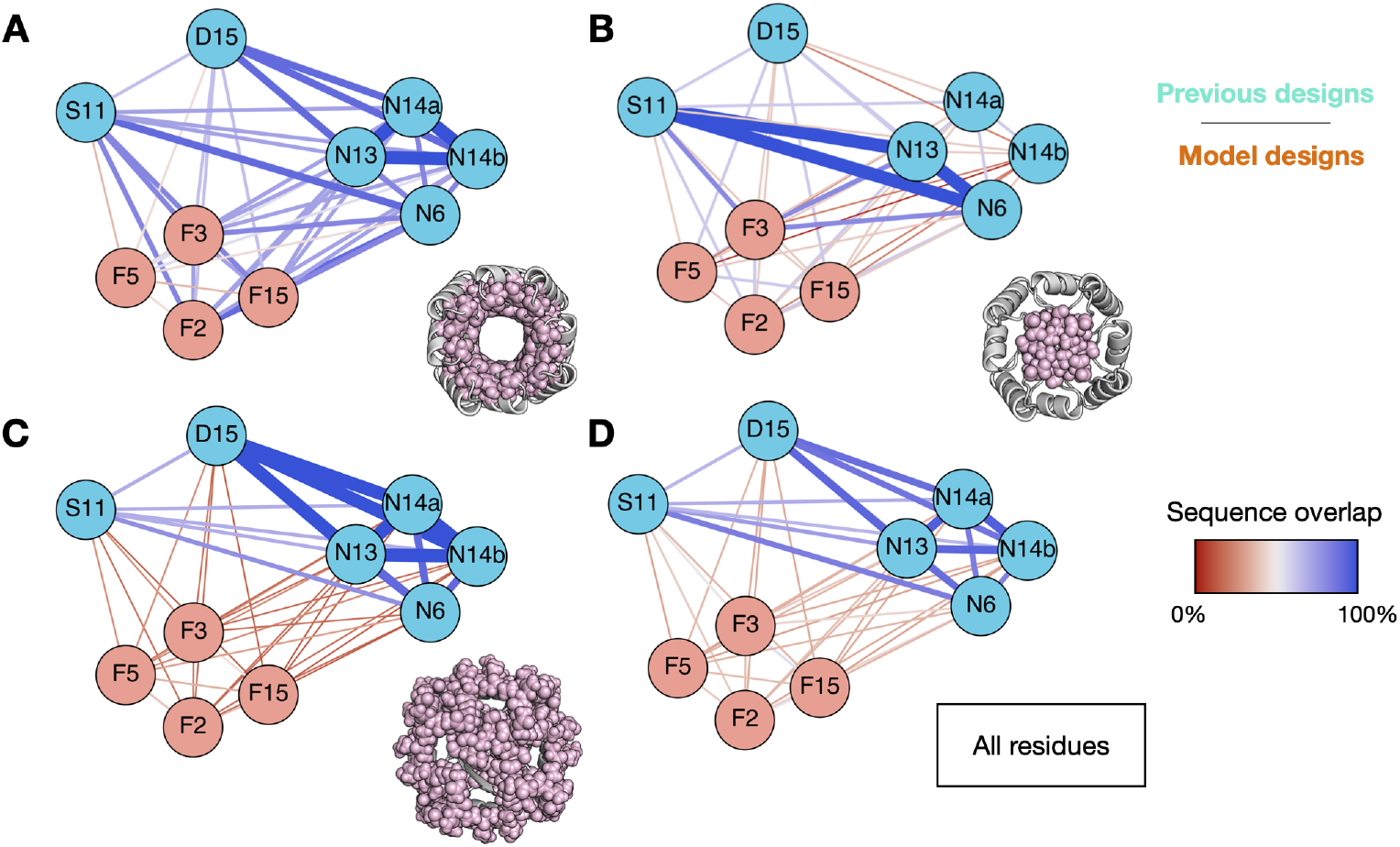
Sequence overlap between TIM-barrel subunits for model TIM-barrel designs (orange) and previously characterized sequences for the same scaffold (blue), including sTIM-11 *(5bvl,* S11) [40], DeNovoTIM15 (6wvs, D15) [44], and NovoTIMs (N6, N13, N14a, N14b) [45]. Specific overlaps shown for the region between (**A**) the helices and the barrel, (**B**) the inner part of the barrel, (**C**) exposed regions, and (**D**) all residues.

**Figure S18:**
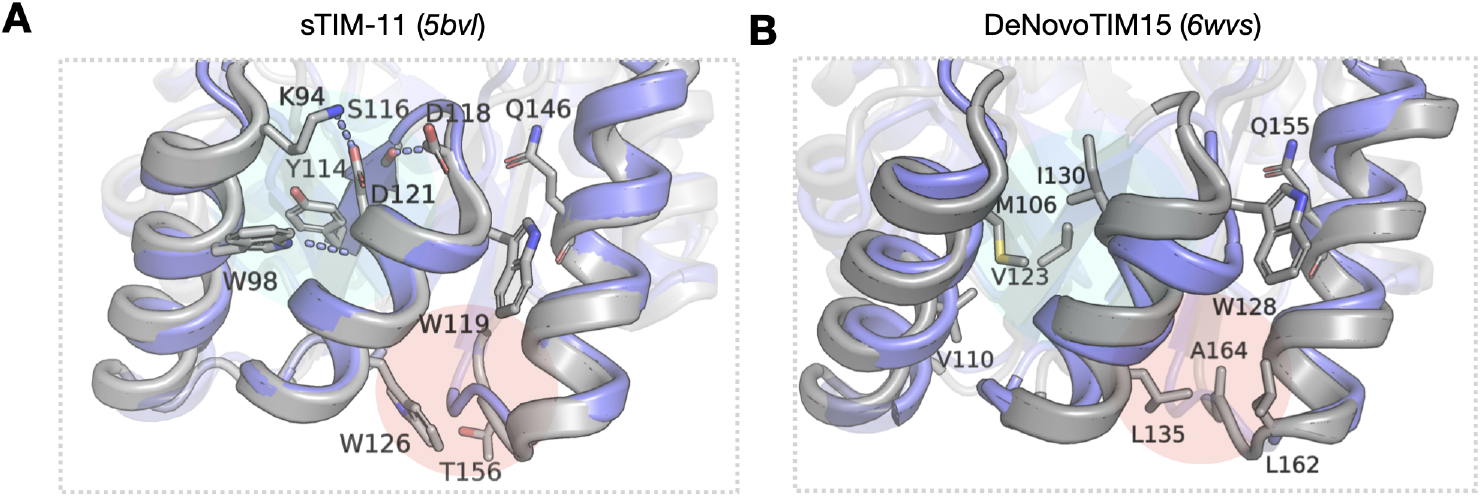
Sequence features for the symmetric subunit near the top of the TIM-barrel (cyan) and the helix interface between symmetric subunits (orange) for previously characterized designs (**A**) sTIM-11 and (**B**) DeNovoTIM15 with sTIM-11 Rosetta model (design scaffold) overlaid in purple.

**Table S1:** Selection of native test case designs for further characterization. Unless otherwise stated, designs were selected based on model PLL rank after applying specified filters across 50 designs. Designs were done on crystal structures as well as crystal structures relaxed under the Rosetta energy function but constrained to the starting coordinates (constrained relaxed backbones). Top sequences highlighted in main paper in Fig. 2I-J are *1acfd3,1bkr d2,1cc8 d2,* and *3mx7 d4*.

| Backbone | Design | Filters and selection procedure |
| --- | --- | --- |
| 1cc8 | Filters | capping residue percentage -- 100%; packstat $\geq 0.55$; packstat post-RosettaRelax $\geq 0.55$ |
|  | d1 | 3rd lowest model negative PLL on constrained relaxed backbone |
|  | d2 | lowest model negative PLL on constrained relaxed backbone |
|  | d3 | lowest model negative PLL on crystal structure |
|  | d4 | 2nd lowest model negative PLL on constrained relaxed backbone |
| 3mx7 | Filters | packstat $\geq 0.55$; packstat post-RosettaRelax $\geq 0.55$; sidechain buried unsats $\leq 2$ |
|  | d1 | 2nd lowest model negative PLL on constrained relaxed backbone |
|  | d2 | lowest model negative PLL on constrained relaxed backbone |
|  | d3 | lowest model negative PLL on crystal structure |
|  | d4 | 2nd lowest model negative PLL on crystal structure |
| 1acf | Filters | packstat $\geq 0.55$; packstat post-RosettaRelax $\geq 0.55$; backbone buried unsats $\leq 10$ |
|  | d1 | ranked 15th in model PLL on constrained relaxed backbone. Has designed disulfide. Negative control for PLL ranking. |
|  | d2 | lowest model negative PLL on constrained relaxed backbone without packstat filters (packstat -- 0.54, 0.487 post RosettaRelax) |
|  | d3 | lowest model negative PLL on crystal structure |
|  | d4 | lowest model negative PLL on constrained relaxed backbone |
| 1bkr | Filters | capping residue percentage -- 100%; packstat $\geq 0.55$; packstat post-RosettaRelax $\geq 0.55$ |
|  | d1 | ranked 12th in model PLL on constrained relaxed backbone. Negative control for PLL ranking. |
|  | d2 | 4th lowest model negative PLL on constrained relaxed backbone and highest post-RosettaRelax packstat among top 4 designs ranked by model PLL |
|  | d3 | 2nd lowest model negative PLL on crystal structure without packstat filters (packstat -- 0.507, 0.562 post RosettaRelax) |
|  | d4 | 3rd lowest model negative PLL on crystal structure and highest post-RosettaRelax packstat among top 3 designs ranked by model PLL |

**Table S2:** TIM-barrel sequence selection – metrics-based selection of 13 designs from initial 50 after filtering by packstat ≥ 0.55 pre- and post-RosettaRelax and 100% occurrence of N-terminal helical capping residues. For each metric, the corresponding rank among 50 designs is given. Metrics used to select a particular sequence are highlighted in red. Secondary metrics of interest are bolded. For instance, F7 has high rank in terms of model PLL but also ranks highly in terms of exposed hydrophobics score. (Blue rows) Crystal structures. (Blue, green rows) Folded designs. (Orange rows) Tested designs that do not appear folded or are not monodisperse based on SEC.

| Design | Sequence rank for metrics of interest across 50 designs |  |  |  |  |  |  |  |  |  |  |  |  |
| --- | --- | --- | --- | --- | --- | --- | --- | --- | --- | --- | --- | --- | --- |
|  | Post-design metrics |  |  |  |  | Post-RosettaRelax metrics |  |  |  |  |  | Psipred cross entropy |  |
|  | model PLL | BB buried unsats | Exposed hydrophobics | Packstat | SC buried unsats | model PLL | BB buried unsats | Exposed hydrophobics | Packstat | CA RMSD | SC buried unsats | MSA | Single seq |
| F1 | 35 | 23 | 28 | 34 | 18 | 2 | 2 | 32 | 2 | 3 | 30 | 18 | 25 |
| F2 | 12 | 5 | 34 | 32 | 7 | 6 | 1 | 34 | 24 | 10 | 30 | 43 | 8 |
| F15 | 5 | 1 | 8 | 24 | 5 | 13 | 40 | 1 | 6 | 45 | 41 | 5 | 37 |
| F3 | 8 | 23 | 27 | 47 | 18 | 18 | 4 | 23 | 45 | 37 | 20 | 19 | 27 |
| F14 | 18 | 3 | 43 | 33 | 23 | 34 | 22 | 38 | 13 | 28 | 1 | 45 | 19 |
| F5 | 23 | 34 | 42 | 46 | 29 | 4 | 30 | 43 | 14 | 13 | 30 | 27 | 16 |
| F4 | 16 | 31 | 47 | 3 | 37 | 31 | 24 | 47 | 20 | 16 | 38 | 1 | 3 |
| F8 | 38 | 5 | 35 | 22 | 45 | 42 | 40 | 35 | 1 | 47 | 40 | 40 | 47 |
| F11 | 46 | 38 | 23 | 43 | 23 | 26 | 47 | 25 | 15 | 8 | 1 | 23 | 42 |
| F6 | 3 | 4 | 4 | 37 | 1 | 20 | 6 | 4 | 21 | 34 | 1 | 4 | 1 |
| F10 | 6 | 27 | 14 | 12 | 38 | 1 | 28 | 18 | 8 | 11 | 38 | 47 | 32 |
| F9 | 22 | 23 | 9 | 2 | 29 | 17 | 15 | 6 | 7 | 41 | 30 | 14 | 7 |
| F7 | 1 | 9 | 4 | 19 | 9 | 32 | 24 | 8 | 31 | 32 | 14 | 20 | 22 |

**Table S3:**
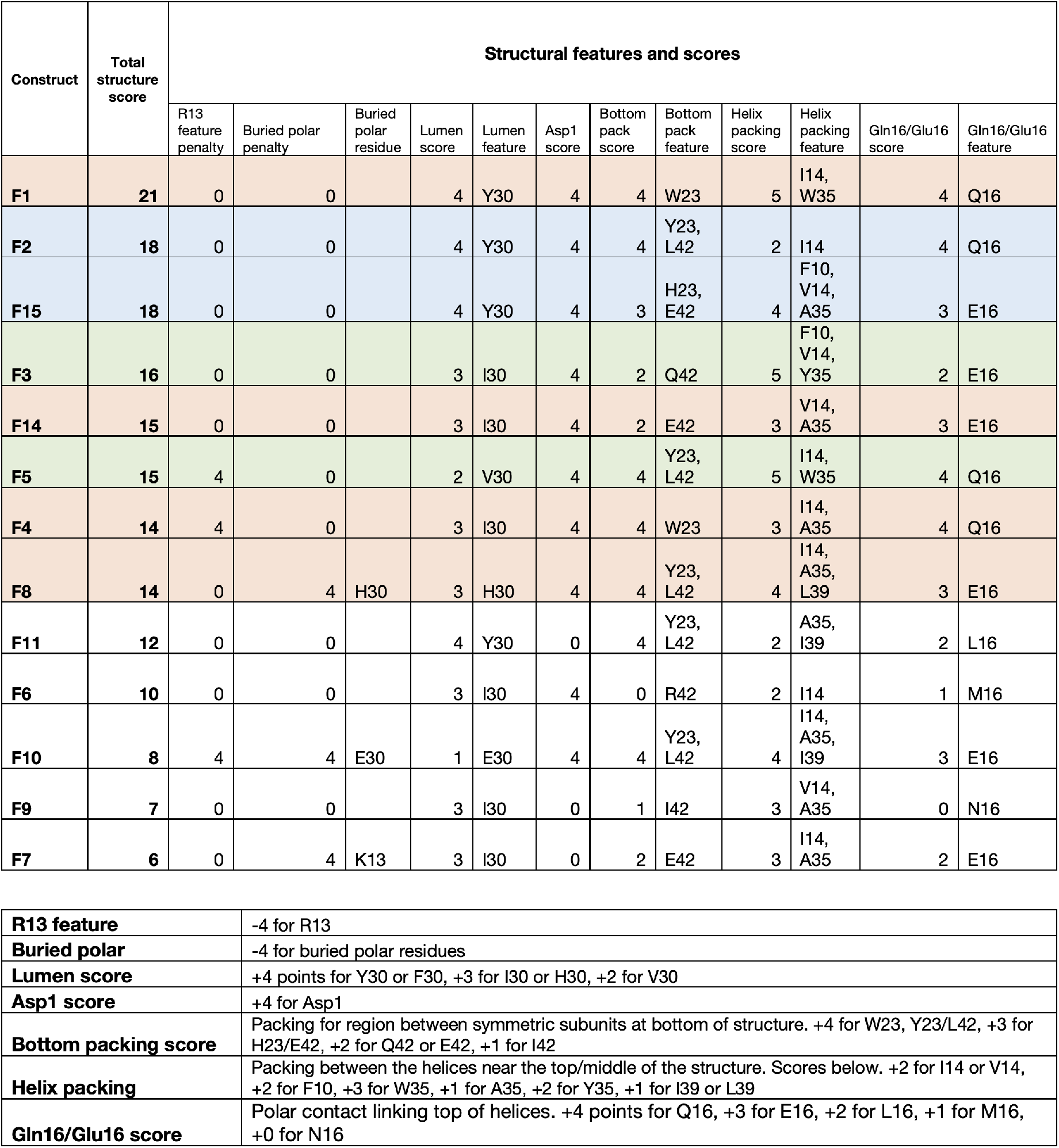
TIM-barrel sequence selection – structural feature-based selection. Selection of top 8 designs from 13 based on simple criteria for a set of structural features. (Top) Total scores, scores across different critera, and corresponding features for criteria. (Bottom) Description of criteria and scoring methodology. (Blue rows) Crystal structures. (Blue, green rows) Folded designs. (Orange rows) Tested designs that do not appear folded or are not monodisperse based on SEC.

| Construct | Total structure score | Structural features and scores |  |  |  |  |  |  |  |  |  |  |  |
| --- | --- | --- | --- | --- | --- | --- | --- | --- | --- | --- | --- | --- | --- |
|  |  | R13 feature penalty | Buried polar penalty | Buried polar residue | Lumen score | Lumen feature | Asp1 score | Bottom pack score | Bottom pack feature | Helix packing score | Helix packing feature | Gln16/Glu16 score | Gln16/Glu16 feature |
| F1 | 21 | 0 | 0 |  | 4 | Y30 | 4 | 4 | W23 | 5 | I14, W35 | 4 | Q16 |
| F2 | 18 | 0 | 0 |  | 4 | Y30 | 4 | 4 | Y23, L42 | 2 | I14 | 4 | Q16 |
| F15 | 18 | 0 | 0 |  | 4 | Y30 | 4 | 3 | H23, E42 | 4 | F10, V14, A35 | 3 | E16 |
| F3 | 16 | 0 | 0 |  | 3 | I30 | 4 | 2 | Q42 | 5 | F10, V14, Y35 | 2 | E16 |
| F14 | 15 | 0 | 0 |  | 3 | I30 | 4 | 2 | E42 | 3 | V14, A35 | 3 | E16 |
| F5 | 15 | 4 | 0 |  | 2 | V30 | 4 | 4 | Y23, L42 | 5 | I14, W35 | 4 | Q16 |
| F4 | 14 | 4 | 0 |  | 3 | I30 | 4 | 4 | W23 | 3 | I14, A35 | 4 | Q16 |
| F8 | 14 | 0 | 4 | H30 | 3 | H30 | 4 | 4 | Y23, L42 | 4 | I14, A35, L39 | 3 | E16 |
| F11 | 12 | 0 | 0 |  | 4 | Y30 | 0 | 4 | Y23, L42 | 2 | A35, I39 | 2 | L16 |
| F6 | 10 | 0 | 0 |  | 3 | I30 | 4 | 0 | R42 | 2 | I14 | 1 | M16 |
| F10 | 8 | 4 | 4 | E30 | 1 | E30 | 4 | 4 | Y23, L42 | 4 | I14, A35, I39 | 3 | E16 |
| F9 | 7 | 0 | 0 |  | 3 | I30 | 0 | 1 | I42 | 3 | V14, A35 | 0 | N16 |
| F7 | 6 | 0 | 4 | K13 | 3 | I30 | 0 | 2 | E42 | 3 | I14, A35 | 2 | E16 |

**Table S3:** TIM-barrel sequence selection – structural feature-based selection.
|  |  |
| --- | --- |
| **R13 feature** | -4 for R13 |
| **Buried polar** | -4 for buried polar residues |
| **Lumen score** | +4 points for Y30 or F30, +3 for I30 or H30, +2 for V30 |
| **Asp1 score** | +4 for Asp1 |
| **Bottom packing score** | Packing for region between symmetric subunits at bottom of structure. +4 for W23, Y23/L42, +3 for H23/E42, +2 for Q42 or E42, +1 for I42 |
| **Helix packing** | Packing between the helices near the top/middle of the structure. Scores below. +2 for I14 or V14, +2 for F10, +3 for W35, +1 for A35, +2 for Y35, +1 for I39 or L39 |
| **Gln16/Glu16 score** | Polar contact linking top of helices. +4 points for Q16, +3 for E16, +2 for L16, +1 for M16, +0 for N16 |

**Table S4:** Sequence alignment of symmetric subunits for previously characterized four-fold or nearly four-fold symmetric TIM barrel designs (sTIM-11, DeNovoTIM15, NovoTIMs) with model designs (FX). Mutations relative to sTIM-11 highlighted in red. Blue columns indicate positions where the previous designs are convergent and one or more model designs differ. Green columns indicate positions where one or more model designs differ from sTIM-11, as does a previously characterized design.

|  | 1 | 2 | 3 | 4 | 5 | 6 | 7 | 8 | 9 | 10 | 11 | 12 | 13 | 14 | 15 | 16 | 17 | 18 | 19 | 20 | 21 | 22 | 23 |
| --- | --- | --- | --- | --- | --- | --- | --- | --- | --- | --- | --- | --- | --- | --- | --- | --- | --- | --- | --- | --- | --- | --- | --- |
| sTIM11 | D | I | L | I | V/C | D | A | T | D | K | D | E | A | W | K | Q/C | V | E | Q | L | R | R | E |
| DeNovoTIM15 | D | I | L | I | V | N | A | T | D | V | D | E | M | L | K | Q | V | E | I | L | R | R | L |
| NovoTIM6 | D | I | L | I | V | D | A | T | D | V | D | E | A | W | K | Q | V | E | I | L | R | R | L |
| NovoTIM13 | D | I | L | I | V | D | A | T | D | V | D | E | M | L | K | Q | V | E | I | L | R | R | L |
| NovoTIM14a | D | I | L | I | V | D | A | T | D | V | D | E | M | L | K | Q | V | E | I | L | R | R | L |
| NovoTIM14b | D | V | L | I | V | D | A | T | D | V | D | E | M | L | K | Q | V | E | I | L | R | R | L |
| F2 | D | I | A | I | V | D | A | D | N | P | A | D | A | I | Q | Q | V | K | D | L | R | K | Y |
| F15 | D | I | L | I | C->V | N | P | D | D | F | E | K | G | V | E | E | V | K | E | L | K | R | H |
| F5 | D | I | V | I | M | N | H | D | D | L | A | E | R | I | K | Q | V | Q | R | L | K | E | Y |
| F3 | D | I | L | I | V | D | H | A | D | F | D | K | A | V | E | E | V | K | Q | L | K | K | E |

**Table S4:** Sequence alignment of symmetric subunits for previously characterized four-fold or nearly four-fold symmetric TIM barrel designs (sTIM-11, DeNovoTIM15, NovoTIMs) with model designs (FX).
|  | 24 | 25 | 26 | 27 | 28 | 29 | 30 | 31 | 32 | 33 | 34 | 35 | 36 | 37 | 38 | 39 | 40 | 41 | 42 | 43 | 44 | 45 | 46 |
| --- | --- | --- | --- | --- | --- | --- | --- | --- | --- | --- | --- | --- | --- | --- | --- | --- | --- | --- | --- | --- | --- | --- | --- |
| sTIM11 | G | A | T | Q | I | A | Y | R | S | D | D | W | R | D | L | K | E | A | W | K | K | G | A |
| DeNovoTIM15 | G | A | K | Q | I | A | V | V | S | D | D | W | R | I | L | Q | E | A | L | K | K | G | G |
| NovoTIM6 | G | A | K | Q | I | A | Y | R | S | D | D | W | R | D | L | Q | E | A | L | K | K | G | A |
| NovoTIM13 | G | A | K | Q | I | A | V | R | S | D | D | W | R | I | L | Q | E | A | L | K | K | G | A |
| NovoTIM14a | G | A | K | Q | I | W | V | I | S | D | D | W | R | I | L | Q | E | A | L | K | K | G | A |
| NovoTIM14b | G | A | K | Q | I | V | V | I | S | D | D | W | R | I | L | Q | E | A | L | K | K | G | A |
| F2 | G | A | K | L | I | A | Y | K | S | K | S | S | E | E | L | K | L | A | L | K | A | G | A |
| F15 | G | A | K | I | I | A | Y | I | S | K | S | A | E | E | L | K | K | A | E | K | A | G | A |
| F5 | G | A | K | Y | I | A | V | L | T | K | N | W | E | D | G | V | K | A | L | K | A | G | A |
| F3 | G | A | K | Y | I | A | I | R | S | D | D | Y | K | K | L | K | K | A | Q | D | A | G | A |

**Table S5:**
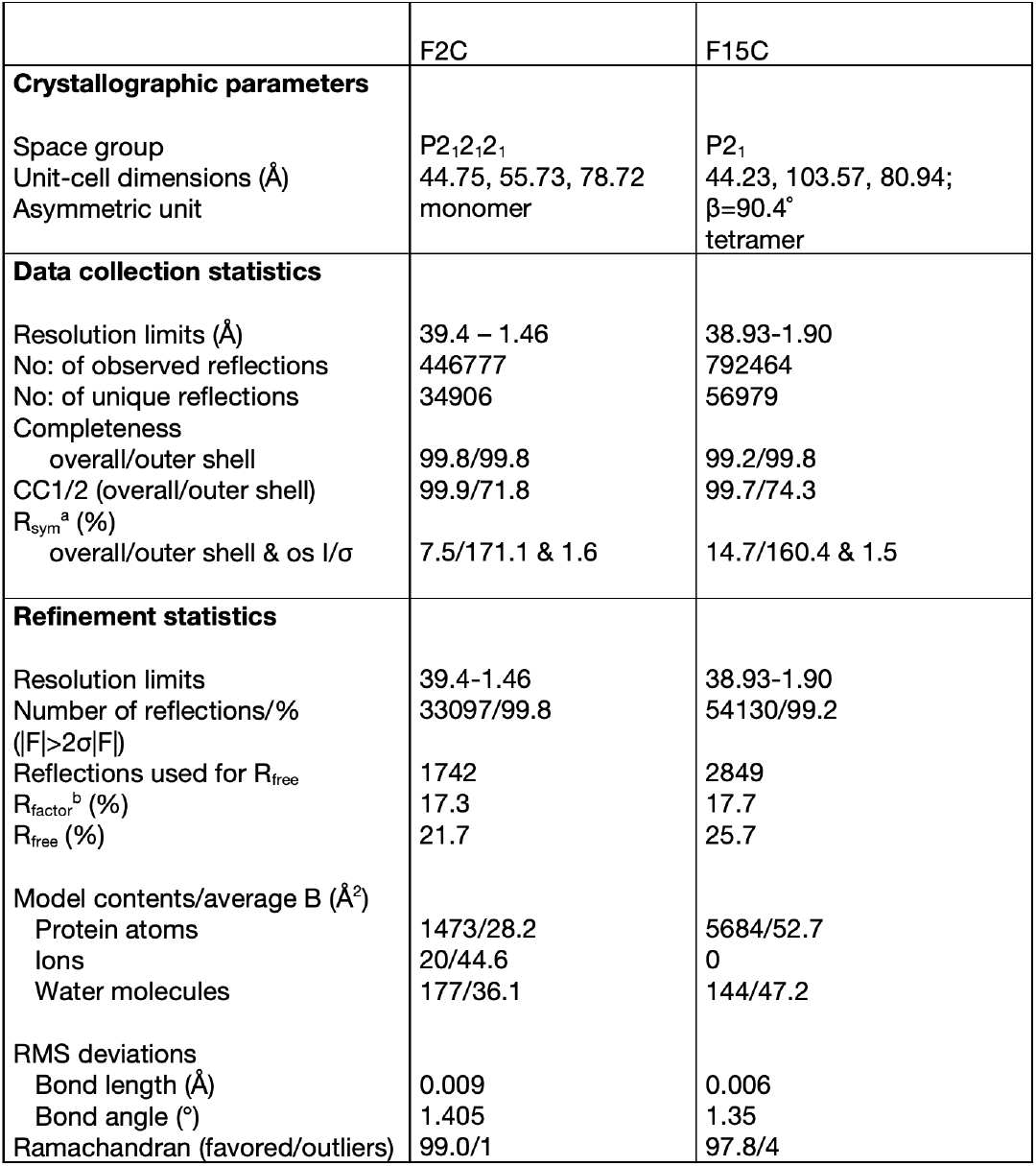
Crystallographic parameters, data collection, and refinement statistics for TIM-barrel crystal structures.

**Table S6:** TIM-barrel crystal structure alpha-carbon (CA) RMSD (Å) to design scaffold prepared with sTIM-11 sequence. CA RMSD for each symmetric unit, all aligned alpha-carbons, and all alpha-carbons. Data for F2C, four F15C monomers, 5bvl [40], and 6wvs [44].

| Structure | Unit 1 | Unit 2 | Unit 3 | Unit 4 | All (aligned) | All |
| --- | --- | --- | --- | --- | --- | --- |
| F2C | 0.829 | 0.7 | 0.617 | 5.958 | 0.769 | 3.346 |
| F15C A | 0.827 | 0.685 | 0.549 | 0.602 | 0.757 | 0.879 |
| F15C B | 0.995 | 0.5 | 2.367 | 1.085 | 0.741 | 1.184 |
| F15C C | 1.607 | 0.575 | 0.591 | 0.931 | 0.878 | 1.27 |
| F15C D | 1.027 | 0.622 | 0.665 | 0.613 | 0.826 | 0.987 |
| 5bvl | 1.164 | 0.779 | 0.746 | 0.831 | 0.679 | 1.256 |
| 6wvs | 1.178 | 1.466 | 1.576 | 1.49 | 1.407 | 1.541 |

